# Serotonergic modulation of visual neurons in *Drosophila melanogaster*

**DOI:** 10.1101/619759

**Authors:** Maureen M Sampson, Katherine M Myers Gschweng, Ben J Hardcastle, Shivan L Bonanno, Tyler R Sizemore, Rebecca C Arnold, Fuying Gao, Andrew M Dacks, Mark A Frye, David E Krantz

## Abstract

Sensory systems rely on neuromodulators, such as serotonin, to provide flexibility for information processing in the face of a highly variable stimulus space. Serotonergic neurons broadly innervate the optic ganglia of *Drosophila melanogaster*, a widely used model for studying vision. The role for serotonergic signaling in the *Drosophila* optic lobe and the mechanisms by which serotonin regulates visual neurons remain unclear. Here we map the expression patterns of serotonin receptors in the visual system, focusing on a subset of cells with processes in the first optic ganglion, the lamina, and show that serotonin can modulate visual responses. Serotonin receptors are expressed in several types of columnar cells in the lamina including 5-HT2B in lamina monopolar cell L2, required for the initial steps of visual processing, and both 5-HT1A and 5-HT1B in T1 cells, whose function is unknown. Subcellular mapping with GFP-tagged 5-HT2B and 5-HT1A constructs indicates that these receptors localize to layer M2 of the medulla, proximal to serotonergic boutons, suggesting that the medulla is the primary site of serotonergic regulation for these neurons. Serotonin increases intracellular calcium in L2 terminals in layer M2 and alters the kinetics of visually induced calcium transients in L2 neurons following dark flashes. These effects were not observed in flies without a functional 5-HT2B, which displayed severe differences in the amplitude and kinetics of their calcium response to both dark and light flashes. While we did not detect serotonin receptor expression in L1 neurons, they also undergo serotonin-induced calcium changes, presumably via cell non-autonomous signaling pathways. We provide the first functional data showing a role for serotonergic neuromodulation of neurons required for initiating visual processing in *Drosophila* and establish a new platform for investigating the serotonergic neuromodulation of sensory networks.

**Author Summary:** Serotonergic neurons innervate the *Drosophila melanogaster* eye, but the function of serotonergic signaling is not known. We found that serotonin receptors are expressed in all neuropils of the optic lobe and identify specific neurons involved in visual information processing that express serotonin receptors. We then demonstrate that activation of these receptors can alter how visual information is processed. These are the first data suggesting a functional role for serotonergic signaling in *Drosophila* vision. This study contributes to the understanding of serotonin biology and modulation of sensory circuits.

## Introduction

Serotonin acts a neuromodulator [1-5] in a variety of networks including the sensory systems required for olfaction, hearing, and vision [6-17]. In the mammalian visual cortex, serotonin regulates the balance of excitation and inhibition [6], cellular plasticity [18-21], and response gain [8, 22]. In some cases, the contribution of individual receptor subtypes is known; for example, in the mammalian retina, serotonergic signaling reduces GABAergic amacrine cell input to retinal ganglion cells via 5-HT1A [23] and can modulate the response to visual stimuli [24]. However, for most sensory circuits, the manner in which serotonin receptor activation is integrated to regulate information processing remains poorly understood.

The visual system of *Drosophila melanogaster* provides a powerful genetic model to study the mechanisms underlying visual circuit activity and regulation [25, 26]. In *Drosophila,* visual processing begins in the lamina where intrinsic monopolar neurons receive direct input from photoreceptors [27]. Lamina monopolar cells L1 and L2 are first-order neurons that feed into pathways discriminating light “ON” (i.e., increase in luminance) and light “OFF” (i.e., decrease in luminance) stimuli respectively [28, 29]. L1 and L2 neurons respond to changes in luminance in a physiologically identical manner [30-32], while downstream neurons in the medulla transform this information to discriminate ON versus OFF stimuli [29]. Further processing occurs in the lobula and lobula plate to mediate higher-order computations for both motion and contrast detection [29, 33, 34]. Significant progress has been made in mapping the synaptic connectivity and function of visual processing neurons, including those required for motion detection [27, 35-40].

Additional studies have shown that aminergic neuromodulators can modulate visual information processing in flies and other insects [26, 41-46]. Octopamine, the invertebrate equivalent of noradrenaline, is present in processes innervating the medulla and lobula in *Drosophila*, where it regulates state-dependent modulation of visual interneurons [42, 46] including the saliency of objects during flight [43]. Serotonergic neurons also innervate the optic ganglia [47-52] and previous studies indicate that serotonin has an impact on both cellular activity and visual behavior in insects [44, 45, 53, 54]. In the blowfly, serotonin alters electrophysiological field recordings representing the combined output of lamina neurons [53]. In the honeybee, single cell recordings in motion-sensitive lobula neurons showed that serotonergic signaling reduces background activity, directional selectivity, and the amplitude of field potentials evoked by moving stripes [45]. In the house fly, serotonin and other neurotransmitters regulate rhythmic size changes in L1 and L2 terminals in the medulla [55] and in *Drosophila*, serotonin was shown to modulate the voltage dependence of potassium channels in photoreceptors [44]. Despite extensive evidence that serotonin modulates the insect visual system, the molecular mechanisms responsible for these effects and the potential contributions of specific subtypes of serotonin receptors remain unclear.

Serotonin receptor signaling occurs via diverse secondary messenger cascades [56, 57] and receptors may act individually or in combination within a single cell [58, 59] or circuit [60]. Moreover, receptors can have different functions in different cellular compartments and can induce both immediate and long-term changes in cell physiology. In addition, since serotonergic modulation targets cells within larger circuits, it is likely that cell non-autonomous mechanisms can induce regulatory changes in neurons that do not themselves express any serotonin receptors. In sum, the precise effects of serotonin receptor activation in specific cells are difficult to predict and necessitate the use of functional assays to complement expression studies. Although previous studies suggest that serotonin affects visual processing, and two recent studies have reported serotonin receptor expression in visual neurons [61, 62], to our knowledge there is no information on the functional effect(s) of any serotonin receptor in any cell within the insect visual system. In this work, we show how a specific serotonin receptor, 5-HT2B, regulates visual responses in L2 lamina monopolar cells, which are critical for the initiation of visual information processing. Expansion of the approach demonstrated here to other receptors and cells may be used to generate a comprehensive picture of serotonergic neuromodulation within a well-defined sensory circuit.

## Results

### Distinct lamina neurons express different serotonin receptors

Five genes encoding serotonin receptors have been identified in the *Drosophila* genome: 5-HT1A, 5-HT1B, 5-HT2A, 5-HT2B and 5-HT7 [63-67]. To identify specific optic lobe neurons expressing each receptor, we expressed the marker mCD8::RFP (or GFP) under the control of a recently characterized panel of T2A-GAL4 insertions in Minos-Mediated Integration Cassettes (MiMICs) located in serotonin receptor gene introns [68]. The GAL4 sequence was inserted into receptor-encoding genes where it acts as an artificial exon and is expected to “mimic” the endogenous gene expression patterns [69]. Ribosome skipping via T2A allows GAL4 to be expressed as a separate protein, rather than a fusion protein with the serotonin receptor [68, 70].

We observed distinct expression patterns for each receptor including projections into the optic lobe neuropils: the lamina (la), medulla (me), lobula (lo) and lobula plate (lp) (S1 Fig). We focus here on receptor subtypes showing expression in the lamina because of the ease of identifying cells based on their morphology [71] as well as the prominent role of this structure in the early stages of visual processing [28, 29, 72]. Since the primary goal of our study was to establish a role in the visual system for one or more of the receptors, it was essential that the function of some identified cells had been previously characterized.

To identify the specific cell types that express each serotonin receptor in the lamina, we used the receptor MiMIC-T2A-GAL4 lines described above in combination with the sparse labeling technique MultiColor FlpOut 1 (MCFO) [73]. Importantly, in contrast to most neurons in the central brain, the stereotyped position and morphology of all neurons in the lamina as well as their organization into repetitive arrays allow them to be identified on the basis of their shape and location alone. Although MCFO can be used for lineage tracing [73], we induced MCFO in adult flies, when visual system neurons are post-mitotic, and MCFO labeling does not represent clonal events. Using 5-HT1A and 5-HT1B MiMIC-T2A-GAL4 lines with MCFO we observed a subtype of cells with a soma in the medulla cortex, a long basket-like projection in the lamina, and a smaller projection in the medulla (Fig 1A-B). This morphology is identical to that of T1 cells and distinct from other cell types in the lamina (Fig 1C) [71]. T1 cells were labeled in 23 of 31 brains (71%) for 5-HT1A and in 10 of 11 brains (91%) for 5-HT1B. On average, we observed thirteen MCFO-labeled T1 cells per individual optic lobe for 5-HT1A and nine T1 cells per optic lobe for 5-HT1B. These data are consistent with the results of recently published studies that used TAPIN-Seq or FACS-SMART-Seq to analyze expression in T1 as well as other cells in the visual system [61, 62] (see S2 Fig for comparison).

**Fig 1.**
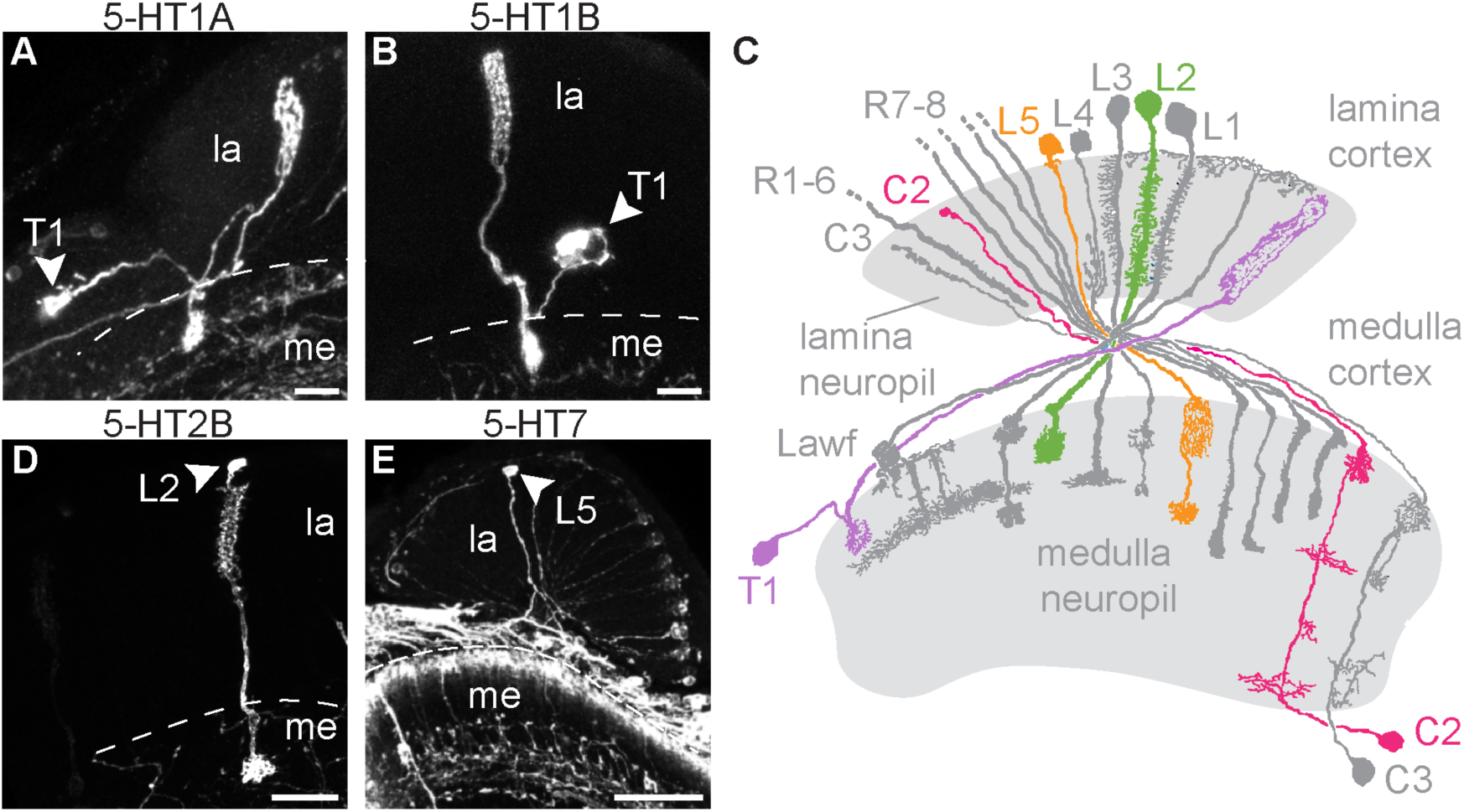
Lamina neurons including T1 and L2 express serotonin receptors. (**A-E**) Serotonin receptor MiMIC-T2A-GAL4 lines were crossed to UAS-MCFO-1 to sparsely label individual cells in the lamina. Cell bodies are indicated by an arrowhead. 5-HT1A (**A**) and 5-HT1B (**B**) MCFO crosses revealed cells with morphologies identical to T1 neurons. (**C**) A diagram showing lamina neurons adapted from [71] highlights L2 (green), L5 (orange), T1 (purple), and C2 (pink). (**D**) 5-HT2B>MCFO labeled cells are morphologically identical to L2 neurons. (**E**) 5-HT7>MCFO-1 labeled neurons most likely representing L5 lamina monopolar cells. Scale bars are 20 μm and n=9-31 brains imaged per receptor subtype. Due to the nature of stochastic labeling, some cell types were observed in only a subset of brains: 22/31 (**A**), 10/11 (**B**), 9/9 (**D**), and 7/13 (**E**).

Using the 5-HT2B MiMIC-T2A-GAL4 driver with MCFO, we observed cells with a soma in the lamina cortex, dense projections extending into the lamina neuropil, and a single bushy terminal in the medulla (Fig 1D), a morphology identical to lamina monopolar neuron L2 and no other lamina cell types (Fig 1C) [71]. We observed L2 cells in 9 of 9 (100%) 5-HT2B>MCFO brains, observing an average of eleven L2 neurons per optic lobe. Additionally, we co-expressed 5-HT2B>RFP with MiMIC-T2A-Lex-ChAT>GFP and found that a subset of the lamina monopolar cells co-labeled with both lines, consistent with the neurochemical identity of L2 cells as cholinergic (S3A Fig, arrowheads). A previous TAPIN-Seq study [61] reported a high probability of 5-HT2B expression in L2 cells, consistent with these findings.

For 5-HT7>MCFO, we observed lamina monopolar cells in 7 of 13 brains (54%) (Fig 1E), with an average of 20 cells per optic lobe. Over 99% of the lamina monopolar cells labeled with 5-HT7>MCFO lacked the dense processes in the neuropil that are characteristic of L1-L3, and also lacked the vertically oriented collaterals in the inner (proximal) lamina seen in L4 neurons (Fig 1C, E). We therefore suggest that 5-HT7 is expressed in the one remaining lamina monopolar cell subtype, L5 cells, consistent with the results of another recent study [61]. Additional examples of 5-HT7>MCFO labeled L5 cells are included in S3B-C Fig.

As noted above, multiple cells of each subtype were generally labeled in each brain. However, we also detected less frequent events for other columnar cells in the lamina. These include observations of multiple centrifugal C2 cells in 3 of 31 5-HT1A>MCFO brains (S3D Fig) consistent with two previous reports of 5-HT1A in C2 [61, 62]. Further supporting the expression of 5-HT1A in C2, we observed co-expression of 5-HT1A>RFP and MiMIC-LexA GAD1>GFP in thin projections in the lamina and in cell bodies between the proximal medulla cortex and the lobula plate (S2E Fig, arrowheads).

### Medulla neurons, glia and serotonergic neurons express serotonin receptors

The results of other more comprehensive studies of RNA expression in visual system neurons prompted us to look beyond our primary focus of lamina neurons [61, 62]. In addition to the columnar neurons that express serotonin receptors and extend projections into the lamina [71], we have tentatively identified cells with projections that are either confined to the medulla, or include both medulla and lobula complex, and may also express serotonin receptors (S3F-H Fig).

For 5-HT2A>MCFO, additional pleomorphic labeling was observed in the lamina cortex (S4A Fig) in a pattern that appeared similar to that of optic lobe glia [74]. Anti-repo labeled nuclei showed close proximity to many 5-HT2A labeled cells (S4B Fig), but one-to-one matching was not possible due to irregular cell morphology. A previous microarray study of glia suggested that 5-HT1A and 5-HT7 are enriched in repo-GAL4 specified glia, while 5-HT1B was enriched in surface glia [75]. Additionally, a previous study [61] reported expression of 5-HT7 in three types of lamina glia—epithelial glia, proximal satellite glia and marginal glia.

The major ganglia of the visual system do not contain any serotonergic cell bodies; rather projections from neurons in the accessory medulla and central brain innervate the optic lobes. Immunolabeling for serotonergic boutons can be observed within all optic ganglia neuropil as well as the lamina cortex (S5A Fig) [50, 76-79]; e.g. arborizations from 5-HT1B>GFP labeled cells in the outer medulla were surrounded by a honeycomb pattern of serotonergic immunolabeling (S5C Fig). Sparse labeling with 5-HT1B>MCFO co-labeled with serotonin immunolabeled boutons in the inner medulla (iM), medulla layer 4 (M4), and lobula (lo) (S5E Fig). Co-labeling with serotonin suggests that 5-HT1B could function as an autoreceptor in these processes. A previous study also reported co-labeling between 5-HT1B-GAL4 and serotonin-immunoreactive cell bodies in the subesophageal ganglion and suggested that 5-HT1B may function as an autoreceptor in serotonergic neurons [80]. Another study sequenced serotonergic neurons specified by TRH-GAL4 and found elevated 5-HT1A, 5-HT1B and 5-HT7 transcripts [81]. Although we did not comprehensively map all putative serotonin autoreceptors in the central brain, we used serotonin cell maps described in [50, 76, 78, 79] to identify 5-HT1B+ cell clusters as LP2 (Cb1), PLP (LP1), and PMP, (S5 Fig) and 5-HT1A+ serotonergic clusters as PLP (LP1), SEL, AMP and PMP (S6 Fig). The integration of functional studies in both serotonergic neurons and post-synaptic neurons represents a future goal to assess the interplay between auto- and post-synaptic receptors in visual circuits.

### L2 neurons express 5-HT2B and T1 neurons express 5-HT1A and 5HT1B

Before embarking on further, time-intensive functional studies of specific neurons in the lamina, we sought to confirm the expression of serotonin receptors independently of both previous studies [61, 62] and the data we obtained using MiMIC-T2A-GAL4 lines. To this end, we used a separate set of split-GAL4 [82] or LexA drivers previously shown to be specific for particular cell types and focused on a small subset of lamina neurons: T1, L1 and L2. Drivers representing each cell were used to express GFP, and the GFP-labeled cells were isolated via Fluorescence Activated Cell Sorting (FACS). RNA was then extracted from T1, L1 and L2 FACS isolates as well as the unlabeled cells. To probe for serotonin receptor expression in each cell type, we used both RNA-Seq (Fig 2 and S1 Table) and RT-qPCR (S7 Fig and S2 Table). For RNA-Seq, we compared the relative abundance for each receptor by calculating Transcripts Per Million (TPMs). We found that 5-HT1A (182±43 TPM±stdev) and 5-HT1B (278±25 TPM±stdev) were more abundant than other serotonin receptors (range 0.1±0.1 to 2.5±4 TPM±stdev) in T1 samples (N=3, Fig 2A and S1 Table). Two previous sequencing studies similarly reported the expression of 5-HT1A and 5-HT1B in T1 [61, 62]. L2 isolates showed higher TPMs for 5-HT2B (130±75 TPM±stdev) compared to other serotonin receptors (range 6±6 to 31±9 TPM±stdev) (Fig 2B and S1 Table), also consistent with the results of another recent transcriptomic study [61]. For RT-qPCR, we calculated enrichment (i.e., fold change) relative to pooled, unlabeled optic lobe cells using the comparative CT method [83]. Relative to unlabeled optic lobe neurons, L2 samples were enriched for 5-HT2B in 4/5 L2 samples but also showed enrichment for 5-HT7 and 5-HT2A in 3/5 and 5-HT2A in 1/5 samples respectively (S7 Fig and S2 Table). Although we cannot rule out the presence of 5-HT2A or 5-HT7 in L2 based on the results of RT-qPCR, our data and those of others [61] suggest that 5-HT2B may be the only serotonin receptor abundantly expressed in L2 cells.

**Fig 2.**
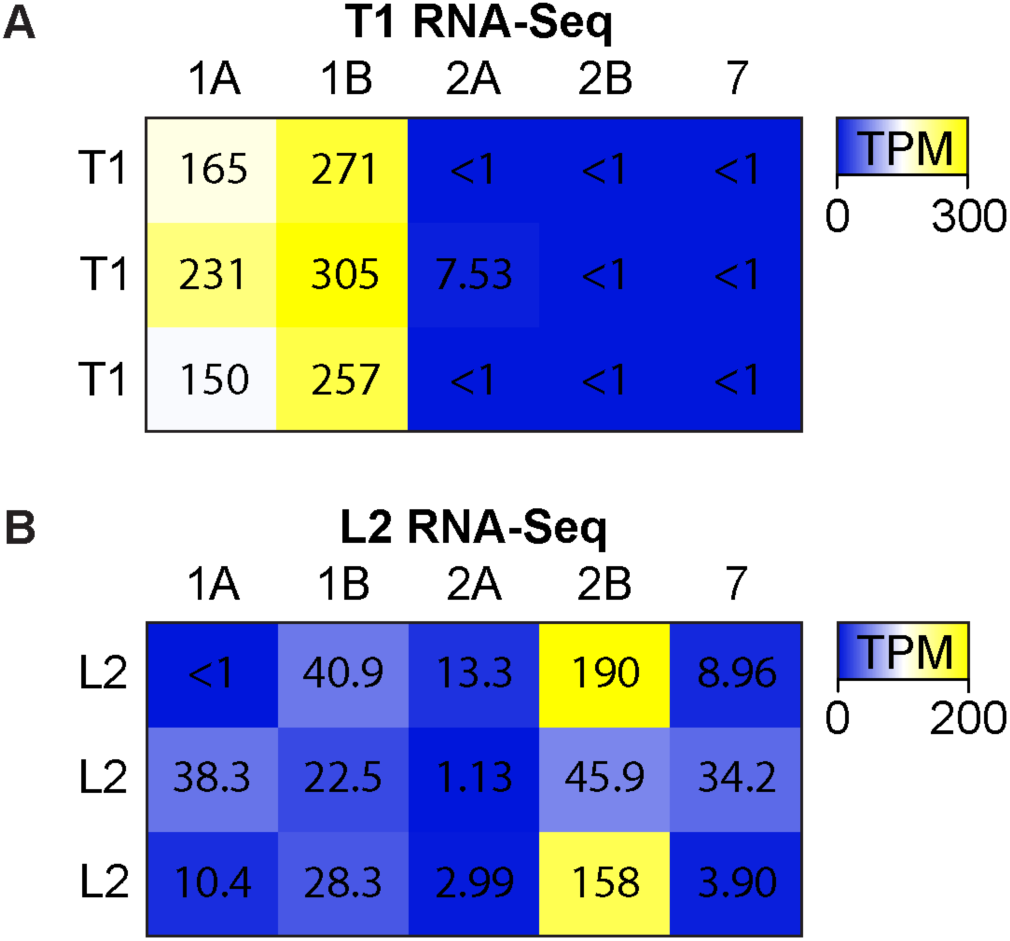
L2 neurons express 5-HT2B and T1 neurons express both 5-HT1A and 5-HT1B serotonin receptors. T1 or L2 neurons were isolated by FACS for RNA-Seq. Transcript abundance was calculated as Transcripts Per Kilobase Million (TPMs). (**A**) T1 RNA-Seq TPMs (color low to high, blue to yellow) were higher for 5-HT1A and 5-HT1B compared to other serotonin receptors. (**C**) L2 RNA-Seq showed high abundance 5-HT2B in two of three samples, with 5-HT2B being the most abundant serotonin receptor transcript in all replicates.

We did not observe evidence of any serotonin receptor expression in L1 neurons using the serotonin receptor MiMIC-T2A-GAL4 lines to drive MCFO. In agreement with this observation, RT-qPCR from isolated L1 cells (S7 Fig and S1 Table) showed virtually no receptor expression, apart from one sample weakly enriched for 5-HT1B. Others have also reported a low likelihood of any serotonin receptor expression in L1 neurons [61]. In sum, MCFO sparse labeling in combination with RNA-Seq and RT-qPCR by our group and others [61, 62] indicate that T1 neurons express 5-HT1A and 5-HT1B, and L2 neurons express 5-HT2B, whereas L1 neurons may not express any serotonin receptor subtypes.

### 5-HT2B and 5-HT1A receptors localize to the medulla neuropil

Both T1 and L2 neurons have dense projections in the lamina neuropil and arborize in layer 2 (M2) of the medulla neuropil. Serotonergic neurons directly innervate M1 and M2 of the medulla neuropil raising the possibility that serotonergic signaling might occur at this site. If so, we reasoned that the serotonin receptors expressed in L2 and T1 might localize to M1 and/or M2. To test this possibility, we took advantage of a 5-HT1A allele that had been tagged at the C-terminus with GFP [84]. The tag was inserted into the endogenous 5-HT1A gene, such that, similar to MiMIC-T2A-GAL4 lines [68, 69], the receptor::GFP fusion protein product is putatively expressed at the same level and in the same cells as the endogenous protein [84]. In 5-HT1A::GFP flies, we observed enrichment of the tagged protein in layer M2 of the medulla relative to other subcellular sites (Fig 3A-A”), suggesting that serotonergic signaling to 5-HT1A::GFP expressing neurons occurs in this region. This might include T1 as well as other columnar neurons that extend processes into M2. Regardless of cell type, there is very low 5-HT1A::GFP signal in the lamina neuropil (Fig 3A’), suggesting that serotonergic signaling primarily targets 5-HT1A receptors localized to the medulla neuropil. We also observed anti-serotonin immunoreactive puncta that co-labeled with 5-HT1A::GFP in the medulla suggesting that 5-HT1A could act as an autoreceptor in serotonergic projection neurons innervating the medulla (Fig 3B, arrows, and S6 Fig).

**Fig 3.**
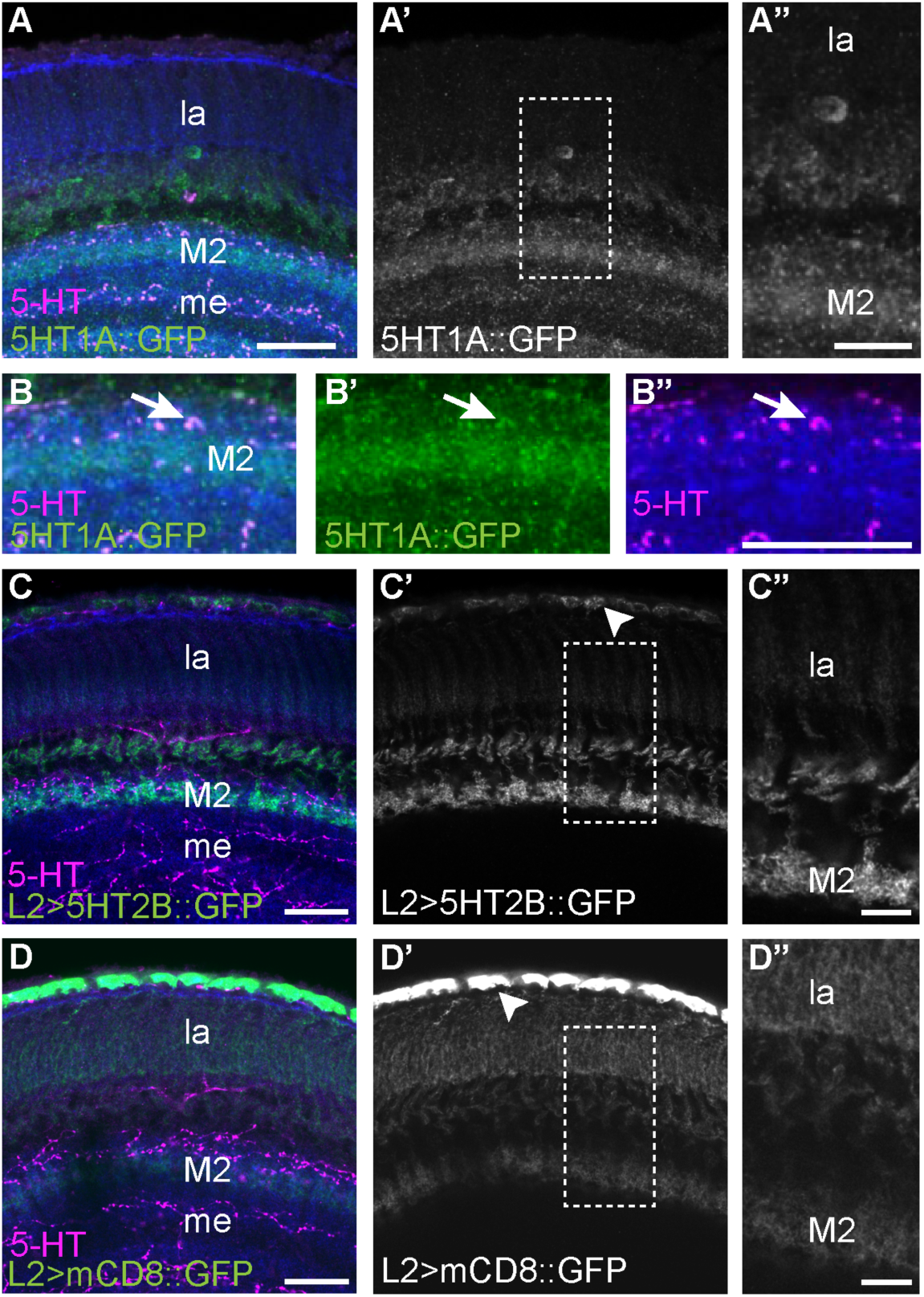
Serotonin receptors 5-HT1A and 5-HT2B are enriched in layer M2 of the medulla. (**A-A”**) 5-HT1A::GFP localized to layer M2 of the medulla, adjacent to some of the boutons in the medulla neuropil immunogenic for serotonin. Neuropil (anti-N-Cadherin, blue) and serotonin (magenta) labeling provide anatomical context for the lamina (la) and medulla (me). The region specified by the dotted lines in (**A’**) is enlarged in (**A”**). (**B-B”**) In some cases, serotonin immunoreactive boutons appear to be co-labeled rather than adjacent to 5-HT1A::GFP (arrow) possibly representing projections from serotonergic neurons in the central brain. (**C-C”**) Subcellular localization of 5-HT2B was visualized by expressing the fusion construct UAS-5-HT2B::sfGFP in L2 neurons specified by L2-split-Gal4. L2>5-HT2B::sfGFP labeling is stronger in the L2 terminals in medulla layer 2 (M2) compared to L2 projections in the lamina (la). Punctate GFP signal is observed in the L2 cell bodies (arrowhead). The region specified by the dotted lines in (**C’**) is enlarged in (**C”**). (**D-D”**) For comparison, membrane-directed UAS-mCD8::GFP was expressed in L2 neurons. L2>mCD8::GFP signal is similar in the medulla and lamina compartments. Strong GFP signal was observed in the cell bodies (arrowhead). The area within the dotted lines in (**D’**) is enlarged in (**D”**). Scale bars are 20 μm in **A-A’**, **B-B”**, **C-C’** and **D-D’**. Scale bars are 10 μm in **A”** and 8 μm in both **C”** and **D”**. Biological replicates are N=6 for 5-HT1A::GFP (**A-B**), N=15 for L2>5-HT2B::GFP (**C-C”**), and N=10 for L2>mCD8::GFP (**D-D”**).

Since we were unable to obtain an endogenously tagged allele of 5-HT2B, we relied on expression of a UAS-5-HT2B::GFP transgene [85] under the control of L2-split-GAL4 to investigate its subcellular localization (Fig 3C). 5-HT2B::GFP was enriched in terminals within M2 of the medulla as compared to the lamina neuropil (Fig 3C-C”) and showed additional punctate labeling in L2 cell bodies (Fig 3C’, arrowhead). To control for the possibility that all membrane-bound proteins might appear to be enriched in M2, we expressed the plasma membrane marker UAS-mCD8::GFP using the same L2 split-GAL4 driver. In contrast to 5-HT2B::GFP, labeling with mCD8::GFP was most prominent in the cell body and proximal processes with progressively weaker labeling through the lamina and medulla neuropil and no enrichment in layer M2 (Fig 3D”). These data strongly suggest that both 5-HT2B and 5-HT1A preferentially localize to the terminals of L2 and T1 respectively in the medulla layer M2 rather than the lamina neuropil. Serotonergic boutons also localize to several layers within the medulla neuropil but are not found in the lamina neuropil (see S5A). It is therefore more likely that T1 and L2 neurons receive serotonergic signals in the medulla rather than the lamina. This may occur in M2, although we cannot rule out other sites in the medulla where serotonergic boutons are present, but the receptors are less enriched.

To further explore serotonergic signaling to L2 and T1 in the medulla, we used sybGRASP to probe for potential synaptic connections between serotonergic boutons in M2 and the terminals of L2 and T1 [86]. We used the previously established ultrastructural connectivity of L2 onto T1 neurons in the medulla [37, 38] as a positive control to validate the use of sybGRASP in detecting interactions within M2, and obtained a robust signal (S8A Fig). By contrast, we failed to detect a signal in M2 in sybGRASP experiments in which the serotonergic neurons were “presynaptic” to L2, T1 or L1 (S8B-E Fig). These data suggest that serotonergic signaling to L2, T1 (and perhaps other columnar neurons), is more likely to be mediated by volume transmission rather than true synaptic transmission, consistent with the use of volume transmission by most aminergic synapses in mammalian systems [87-90].

### Serotonin increases calcium levels in L2 and L1 neurons

The data presented here and by others [61, 62] strongly suggest that L2 and other cells in the visual system express serotonin receptors but do not address their function. To address the potential effects of serotonin on L2 neurons, we bath applied serotonin to the optic lobe and used live imaging to monitor cellular activity. The data for receptor expression in L2 was strongest for 5-HT2B receptors, which couple with G_q/11_ to increase intracellular calcium *in vitro* [67, 91]. We therefore used the genetically encoded calcium indicator GCaMP6f [92] to follow changes in L2 activity that might be induced by serotonin. We again employed the L2split-GAL4 driver used for transcriptional analysis (Fig 2) to specifically express GCaMP6f in L2 neurons (Fig 4A). Since we observed enrichment of 5-HT2B::sfGFP in L2 terminals in M2, we focused our recordings of calcium signaling on these sites. For each experiment, we first recorded a baseline while perfusing the tissue with saline; the perfusion solution was then switched to either saline containing 100 μM serotonin or saline alone. We included tetrodotoxin (TTX) in the perfusion solution in these experiments to reduce inputs to L2 neurons. TTX has been used by others to study the effects of serotonin in *Drosophila,* and represents a standard method to reduce cell-non autonomous neuronal inputs [93-95]

**Fig 4.**
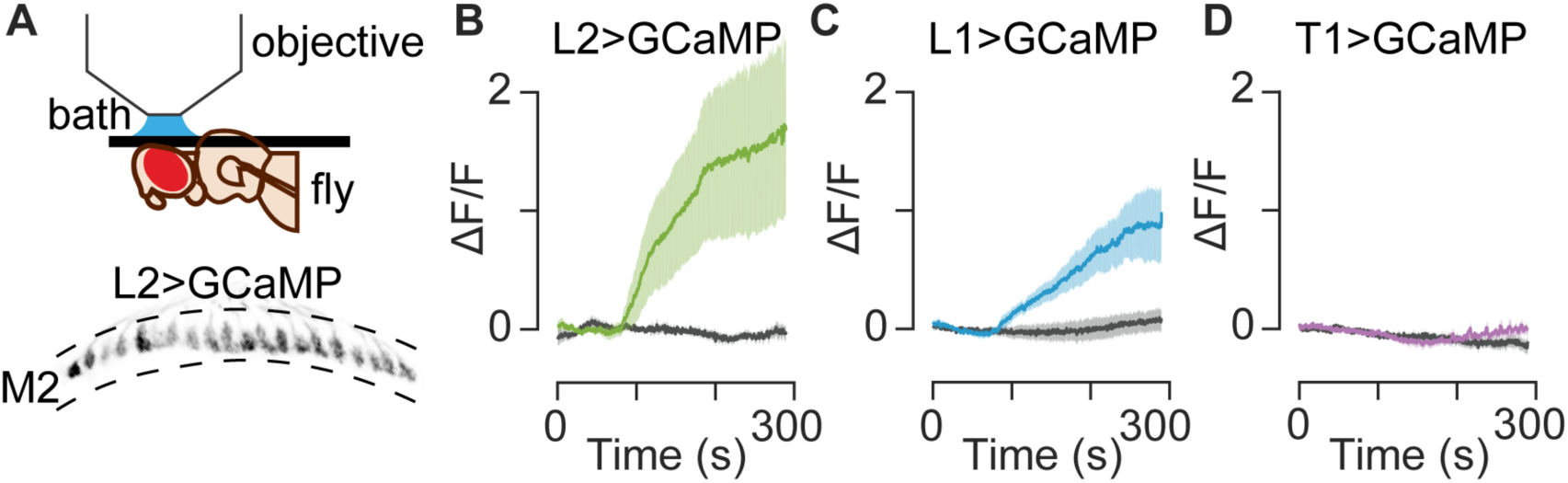
Bath application of serotonin leads to increased calcium in L2 and L1 neurons, but not T1 neurons. (**A**) The experimental setup is shown in the top of panel A, along with a sample image of L2 terminals (bottom of panel A, gray) as imaged in the medulla. (**B-D**) GCaMP6f was paired with L2, L1 and T1 split-GAL4 drivers to monitor responses to 100 μM serotonin (colored traces) or saline controls (gray traces). The perfusion change occurred approximately 105 s into the recording, which corresponds to 45 s in (B-D) because the first 60 s are not shown. (**B**) In L2 terminals serotonin application led to a significant increase in GCaMP6f signal indicating increased calcium levels as compared to saline controls (p=00095). (**C**) L1 terminals showed a similar increase in calcium following a switch to serotonin (p=0.02). (**D**) T1 cells expressing GCaMP6f showed no significant change in calcium following serotonin application (p>0.05). For (B-D) N=4-8 individual flies; the dark trace is an average of all traces and the shaded region is 1 SD; saline vs. serotonin comparisons are two-tailed Wilcoxon rank sum tests.

We consistently observed a large increase in GCaMP6f fluorescence in L2 terminals following serotonin application (Fig 4B and S9A Fig). This increase continued throughout the time course of the recordings, peaking at 1.73 ΔF/F ± 0.77 SEM (compared to saline control - 0.03 ΔF/F ± 0.05 SEM at the same timepoint; p=0.0095 by two-tailed Wilcoxon rank sum test). Thus, serotonin leads to an accumulation of cytosolic calcium in L2 cells, consistent with the predicted outcome of activating G_q/11_ coupled 5-HT2B receptors [67, 91]. Importantly, many neurons in the optic lobe use graded electrical signaling. Synaptic inputs to L2 neurons from these or other cells relatively unaffected by TTX in these experiments could mediate cell non-autonomous effects of serotonin in L2 neurons.

Since we did not detect serotonin receptors in L1 neurons, we did not expect serotonin to measurably change intracellular calcium levels in these cells. However, with GCaMP6f expressed in L1 cells using a cell-specific driver, we regularly observed a robust increase in baseline fluorescence following serotonin exposure (Fig 4C and S9B Fig). Although the increase in GCaMP6f signal did not reach the same response amplitude as observed in L2 neurons, the time course was similar: the signal persisted throughout the recording and peaked at 0.98 ΔF/F ± 0.34 SEM (compared to saline control at 0.07 ΔF/F ± 0.09 SEM; p=0.02 by two-tailed Wilcoxon rank sum test). A direct action of serotonin on L1 is unlikely since neither we (S7C Fig) nor others [61] detect endogenous serotonin receptors in L1 neurons. Possible mechanisms include cell-non-autonomous inputs from either columnar or non-columnar neurons intrinsic to the visual system, more distal projections from the central brain or perhaps electrical coupling between L1 and L2 [28]. The cell non-autonomous inputs mediating the indirect effects of serotonin would seem be more likely to use graded potentials, as is common in the visual system [96-98], since TTX was included in the perfusion solution in these experiments and TTX blocks sodium channels that are required for action potentials [94, 95]. However, we cannot rule out the possibility that TTX blocked channels mediating additional modes of communication.

We next examined whether serotonin could affect the activity of T1 cells. Both 5-HT1A and 5-HT1B receptors, expressed in T1 neurons, are expected to couple with G_i_ proteins and negatively regulate adenylyl cyclase [65, 91]. Due to the generally inhibitory function of these receptors, we hypothesized that serotonin would dampen activity in T1 neurons, possibly manifested as a decrease in cytosolic calcium or membrane potential [99]. Using the T1 split-GAL4 driver [82] to express either GCaMP6f or the voltage sensor Arclight [100], we did not observe a significant change in fluorescence during perfusion with serotonin (Fig 4D, S9C-D Fig) p>0.1). Thus, further experiments will be needed to determine the effects of serotonin on T1 neurons. These negative data are nonetheless important for the current study, since the absence of a GCaMP6f response in T1 neurons indicates that the responses observed in L1 and L2 are not artifacts or a generalized phenomenon common to all cells in the lamina.

### Serotonin in visual processing

To explore the possibility that serotonergic neuromodulation plays a role in visual processing, we tested whether exogenous serotonin alters visually induced calcium transients in L2 neurons. We used GCaMP6f to record and compare calcium transients in flies receiving saline or serotonin perfusion. Previous studies using voltage indicators found that L2 neurons depolarize in response to dark flashes and hyperpolarize in response to light flashes [31]. Similarly, calcium-indicator recordings showed that intracellular calcium increased in the dark and decreased in the light in [30]. Brief light or dark flashes induce bi-phasic calcium transients [31] that enable analysis of calcium kinetics. For this reason, we used brief dark or light flashes to test whether serotonin might alter the magnitude or kinetics of visually induced calcium transients in L2 terminals. Flies were suspended over an LED arena (see Fig 4A) and either a light or dark flash of the entire LED screen (100 ms) was presented at 5 s intervals. Between each flash, the screen showed an intermediate brightness level, indicated as grey in Fig 5A. One-minute “epochs” consisting of 12 flashes of randomly shuffled polarity were presented six times for each trial (Fig 5A). The first 60 s epoch was recorded in saline alone, followed by a switch to either saline with 100 μM serotonin or saline alone during epoch 2 (Fig 5A). Unlike the experiments shown in Fig 4, we did not include TTX in the perfusion solutions because we did not want to affect neuronal communication in any way that might interfere with the response to visual stimuli. Performing additional further experiments in the absence of TTX was also important to rule out the possibility TTX was responsible for the effects seen in Fig 4.

**Fig 5.**
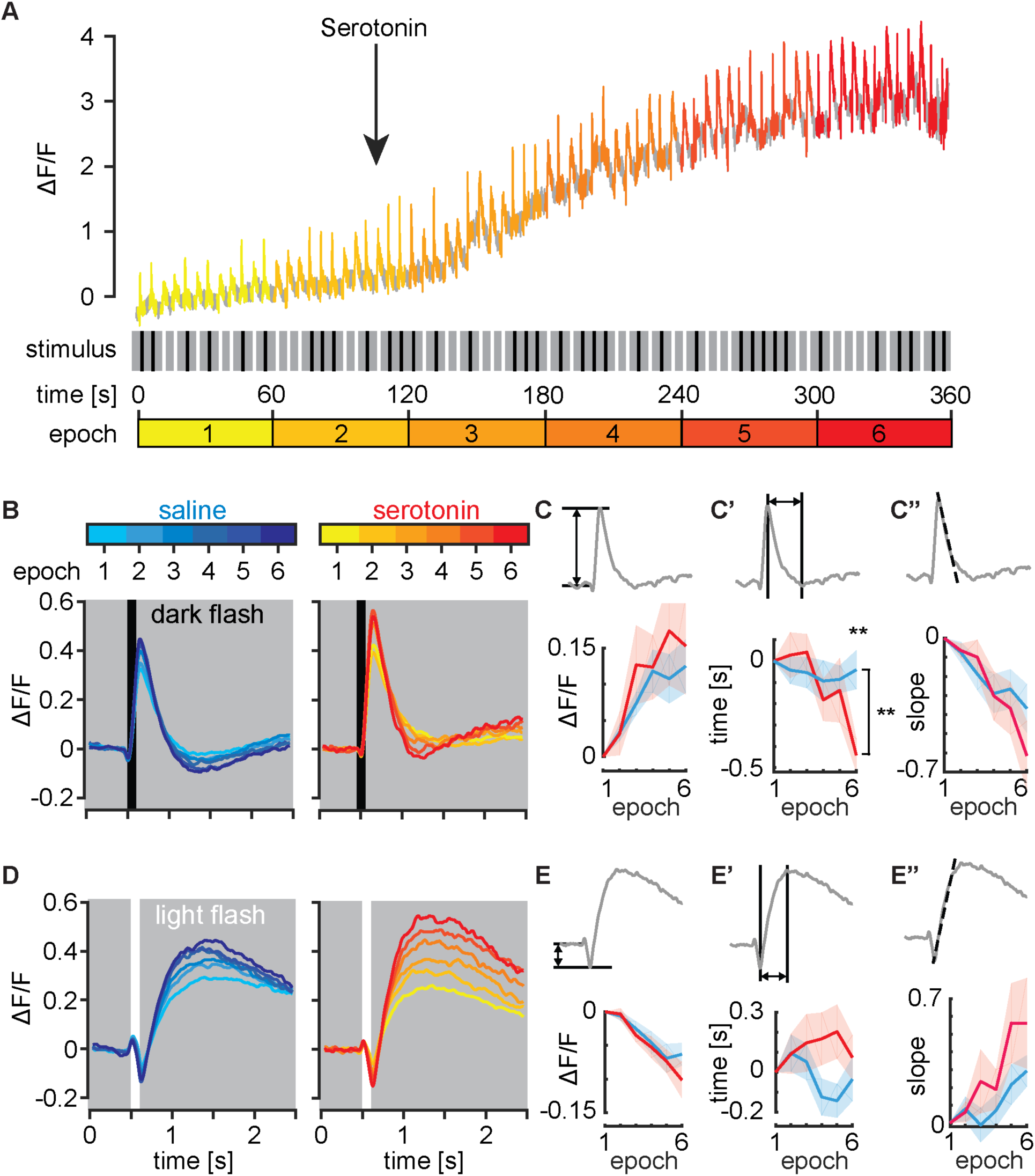
Serotonin modulates L2 neuron visually induced calcium transient kinetics. L2-split-GAL4 was crossed to UAS-GCaMP6f to monitor calcium transients following 100-ms light or dark flashes. The optic lobe was initially perfused with saline alone during a 60 s baseline recording designated as “epoch 1”. The solution was then switched to either saline with serotonin or saline control after 105 s, with the switch occurring during epoch 2. (**A**) A sample recording with serotonin perfusion following the initial baseline is shown in the upper panel. Light or dark stimuli (A, middle panel) were flashed at random every 5 s for 6 min. To visualize changes in the response to light and dark flashes in L2, data from each 60 s epoch was binned (see epochs 1-6 in A, lower panel). (**B**) Response to dark flashes. Color coded traces representing each 60 s epoch are shown for flies receiving saline (left panel, cyan to blue) or serotonin perfusion (right panel, yellow to red). In both groups, L2 terminals responded to a dark flash with a strong increase in GCaMP6f fluorescence before returning to baseline. (**C-C”**) Analysis of dark flash response from panel B. For plotting each variable shown in C, the average value for the epoch 1 baseline was subtracted; epoch 1 is therefore always set to 0 in panel C-C”. (**C**) The change in the amplitude of the calcium transient (the difference between pre-stimulus ΔF/F and maximum ΔF/F) relative to epoch 1 is shown in (C) and did not differ between flies exposed to saline versus serotonin. (**C’**) The change in the time (s) to reach a minimum ΔF/F following the maximum ΔF/F relative to epoch 1 is shown in (C’) and was significantly longer (p≤0.01**) in serotonin-perfused flies compared to saline controls. (**C”**) The change in the slope of the decay from maximum ΔF/F to minimum ΔF/F relative to epoch 1 was not different between the saline and serotonin groups. (**D**) Response to light flashes. When a light flash was presented in control experiments, L2 cells responded with a decrease in GCaMP signal followed by a large sustained rebound. Color coded traces representing each 60 s epoch for saline (left panel, cyan to blue) and serotonin (right panel, yellow to red) are shown. (**E-E”**) Analysis of light flash response from panel D. For plotting each variable shown in E, the average value for the epoch 1 baseline was subtracted; epoch 1 is therefore always set to 0 in panel E-E” as in panel C-C”. (**E**) The change in the amplitude of the initial decrease in calcium (the difference between pre-stimulus ΔF/F and the subsequent minimum ΔF/F) relative to epoch 1 is shown in (E) and was not different between the saline and serotonin groups. (**E’**) The change in the time to reach a maximum ΔF/F (time, s) relative to epoch 1 and (**E”**) the change in the slope of the rebound relative to epoch 1 was not significantly differ between serotonin versus saline controls. Recordings in B-E represent, N=14 and N=20 individual flies perfused with serotonin or saline respectively. Shaded areas show mean +/-SEM. Comparisons are two-way repeated measures ANOVA (brackets show interactions between time and genotype) and Sidak’s multiple comparisons tests, p≤0.05 *, p≤0.01**, p≤0.001***, p≤0.0001****.

Dark flashes induced a large increase in calcium that returned to baseline within ∼1 second as previously described [31] (Fig 5B). The amplitude of the dark flash induced calcium transients increased over the time course of the experiment for animals receiving either serotonin or saline (Fig 5B). To compare differences between the serotonin and saline control groups and the potential effects of serotonin over the time course of the experiment, we quantified three calcium transient variables: amplitude (5C), the time required for decay from the peak to a subsequent minimum (5C’), and the slope of the decay (5C’’). To facilitate the direct comparison of results obtained during perfusion with saline alone versus saline followed by serotonin, we first calculated the average for each variable prior to serotonin exposure measured during epoch-1. We then subtracted the epoch 1 baseline from each epoch, effectively setting the initial value of each plot to 0 for epoch 1 in Fig 5C-C”.

Fig 5C shows the change in calcium transient amplitude, calculated as the difference between the pre-stimulus ΔF/F and the peak ΔF/F, relative to epoch 1. We did not detect a difference between the serotonin and saline groups. Fig 5C’ quantifies the decay time from peak amplitude, calculated as the time (s) between the peak ΔF/F and the subsequent minimum ΔF/F, relative to epoch 1. We detected a modest but statistically significant (p=0.0036 by Repeated Measures ANOVA) increase in the decay time of the dark flash response in animals perfused with serotonin versus saline alone. In Fig 5C” we quantify the change in the slope of the decay relative to epoch 1; we did not detect a difference in slope between flies treated with serotonin versus saline (Fig 5C”).

Light flashes induced a transient decrease in GCaMP6f fluorescence, followed by a secondary sustained rebound (Fig 5D) as previously reported [31]. To analyze calcium transients induced by light flashes, we quantified three variables: the magnitude of the initial calcium decrease relative to the pre-stimulus baseline (Fig 5E), the time from minimal to maximal ΔF/F (Fig 5E’), and the slope of this rebound (Fig 5E’’). Baseline values obtained for the response to light flashes in epoch 1 were subtracted from each epoch to set the initial value of each variable to 0 for epoch 1 in the plots in Fig 5E-E”. We did not observe any significant differences between saline and serotonin groups when calculating the magnitude of the initial downward deflection (Fig 5E), the time from minimal to maximal ΔF/F (Fig 5’) or the slope of the rebound (Fig 5E”).

In sum, serotonin drove a robust increase in intracellular calcium levels of L2 neurons in wildtype flies, regardless of the presence (Fig 4) or absence (Fig 5) of TTX. We also detected a modest, but statistically significant effect of serotonin on the kinetics, but not the amplitude, of the GCaMP6f response of L2 to a dark flash (Fig 5C’). We did not detect any effect of serotonin on the response of L2 to a light flash in 5-HT2B +/+ flies.

### 5-HT2B mediates the effects of serotonin on L2

Since 5-HT2B is expressed in L2 neurons and is predicted to use calcium as a second messenger we hypothesized that it would mediate the response of L2 neurons to serotonin. In flies expressing wild type 5-HT2B we observed a robust serotonin-mediated increase in intracellular calcium (Fig 4B, with TTX in the perfusion solution and Fig 6A, without TTX in the perfusion solution). We next examined whether loss of 5-HT2B would blunt the gradual increase in baseline calcium we observed in flies exposed to serotonin (Fig 6B). As a negative control for experiments using the 5-HT2B homozygous mutants (-/-) we used heterozygous siblings (+/-) in which one wild type allele of 5-HT2B was present (Fig 6B). Similar to 5-HT2B +/+ flies (Fig 6A), 5-HT2B heterozygous controls perfused with serotonin showed a gradual increase in baseline calcium levels over the six-minute time course of the experiment (Fig 6B). In contrast, in the 5-HT2B -/- mutant, the baseline calcium signal was nearly flat over the time course of the experiment (p=0.0024 by two-tailed Wilcoxon rank sum test). To more specifically reduce the expression of 5-HT2B in L2, we also tested available RNAi transgenes directed against 5-HT2B but did not detect a significant effect (data not shown). We therefore cannot rule out indirect cell non-autonomous effects from 5-HT2B expression in other cell types. However, the simplest explanation for the observed results is that activation of 5-HT2B in L2 neurons generates a gradual increase in cytosolic calcium.

**Fig 6.**
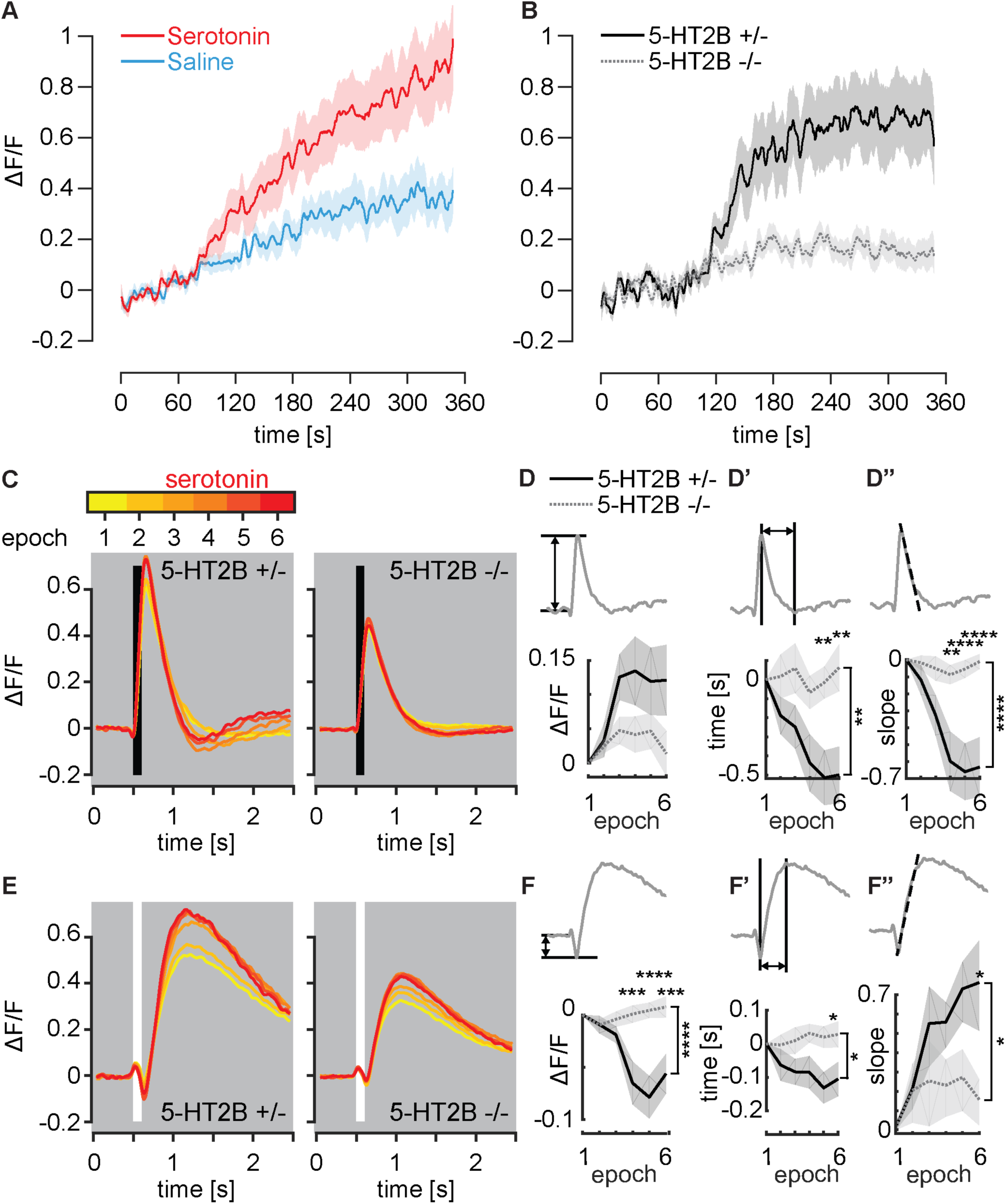
5-HT2B mediates the L2 neuron response to serotonin. (**A**) L2-split-GAL4>GCaMP6f was used to monitor intracellular calcium in L2 terminals. The optic lobe was initially perfused with saline alone and was then switched to either saline with serotonin or saline control after 105 s. Serotonin perfusion (in the absence of TTX) induced an increase in baseline calcium in 5-HT2B +/+ lines receiving serotonin compared to saline controls. (**B**) Mutant 5-HT2B-“KO”-GAL4 flies (5-HT2B -/-) were combined with UAS-GCaMP6f to monitor calcium changes in L2 terminals. Homozygous (5-HT2B -/-) and heterozygous (5-HT2B +/-) flies both received serotonin perfusion. The serotonin-mediated baseline increase was seen in 5-HT2B +/- flies, as in wild type flies, but was not observed in 5-HT2B -/- flies. (**C-F**) Homozygous (5-HT2B -/-) and heterozygous (5-HT2B +/-) 5-HT2B- “KO” -GAL4 flies expressing UAS-GCaMP6f were used to record calcium transients following brief light or dark flashes. The preparation was initially perfused with saline alone during a 60 s baseline recording designated as epoch 1 and the solution was then switched to saline with serotonin after 105 s, with the switch occurring in epoch 2. (**C**) Response to dark flashes. Traces representing each 60 s epoch (yellow to red) are shown for 5-HT2B +/- (left) and 5-HT2B -/- (right). All preparations received serotonin perfusion. In both groups, dark flashes induced a strong, transient increase in GCaMP6f fluorescence that returned to baseline. Dark-flash calcium transient amplitude increased over the course of the experiment for 5-HT2B +/- flies, but this was not seen in 5-HT2B -/- flies. (**D-D”**) Analysis of dark-flash data from panel C. For plotting each variable shown in D-D”, the average value for epoch 1 was subtracted; epoch 1 is therefore always set to 0. (**D**) The change in the amplitude of the maximum calcium transient (the difference between pre-stimulus ΔF/F and maximum ΔF/F) relative to epoch 1 is shown. (**D’**) The change in the time to reach a minimum ΔF/F following the maximum ΔF/F of the calcium transient relative to epoch 1 was significantly decreased in 5-HT2B +/- flies compared to 5-HT2B -/- flies in which this period was essentially unchanged over the time course of the experiment. (**D”**) The change in the slope of the decay relative to epoch 1 was significantly decreased in 5-HT2B +/- flies compared to 5-HT2B -/- flies; it appeared essentially unchanged for 5-HT2B -/- flies over the course of the experiment. (**E**) Response to light flashes. When a light flash was presented, L2 cells responded with a decrease in the GCaMP6f signal followed by a large sustained rebound. Responses for 5-HT2B +/- (left) and 5-HT2B -/- (right) flies are shown. (**F-F”**) Analysis of light flash data from panel E. For plotting each variable shown in F-F”, the average value for the epoch 1 baseline was subtracted; epoch 1 is therefore always set to 0 as in panels D-D”. (**F**) The change in the amplitude of the light flash induced a calcium decrease (calculated as the difference between the pre-stimulus ΔF/F and minimum ΔF/F) relative to epoch 1 that was significantly enhanced over the course of the experiment in 5-HT2B +/- flies relative to 5-HT2B -/- flies. (**F’**) The change in the time to reach a maximum relative to epoch 1 was significantly decreased in 5-HT2B -/- flies compared to the response of 5-HT2B +/- flies which appeared unchanged. (**F”**) The change in the rebound slope relative to epoch 1 increased significantly in 5-HT2B +/- flies relative to 5-HT2B -/- flies. For (A), N=14 serotonin and N=20 saline exposed flies were tested. For (B-F), N=13 5-HT2B +/- and N=15 5-HT2B -/- flies, all receiving serotonin, were tested. Shaded areas show mean +/- SEM. Comparisons in (D, F) are two-way repeated measures ANOVA (brackets show interactions between time and genotype) and Sidak’s multiple comparisons tests, p≤0.05 *, p≤0.01**, p≤0.001***, p≤0.0001****.

We next examined whether loss of 5-HT2B would alter the calcium transients in L2 neuron terminals following light or dark flashes. We hypothesized that loss of 5-HT2B would block any changes in the kinetics of the calcium response seen with serotonin perfusion (see Fig 5C’). We also reasoned that a mutation in the serotonergic signaling pathway might provide a “sensitized genetic background” to enhance the detection of serotonin’s functional effects. As a negative control, we used heterozygous siblings (5-HT2B +/-). We again graphed averages for each epoch after subtraction of the average for epoch-1. For dark flashes, we plotted the change in amplitude of the increase from pre-stimulus ΔF/F to the maximum ΔF/F relative to epoch 1 (Fig 6D), the change in the time between the peak ΔF/F and the subsequent minimum ΔF/F relative to epoch 1 (Fig 6D’), and the change in slope of the decay relative to epoch 1 (Fig 6D”). As we observed in 5-HT2B +/+ flies (see Fig 5C’), perfusion of 5-HT2B +/- heterozygous controls with serotonin led to a gradual reduction in the time required for L2 terminals to reach a minimum after responding to a dark flash (Fig 6D’). In striking contrast, in 5-HT2B -/- homozygous loss of function flies, the time between the peak and the subsequent minimum (Fig 6D’), and the slope of the decay (Fig 6D”) remained unchanged over the course of the experiment and continuous perfusion with serotonin. In sum, the 5-HT2B mutation ablated the serotonin-dependent change in the kinetics of the L2 response to dark flashes. The effects on amplitude (Fig 6D) were not statistically significant between the 5-HT2B -/- mutant and heterozygous controls.

To compare how 5-HT2B +/- and 5-HT2B -/- flies responded to light flashes, we calculated the change in the amplitude of the initial downward deflection from the pre-stimulus baseline relative to epoch 1 (Fig 6F), the change in the time from minimal to maximal ΔF/F relative to epoch 1 (Fig 6F’) and the change in the slope of the rebound relative to epoch 1 (Fig 6F”). The 5-HT2B +/- heterozygotes preparations showed a progressive increase in the magnitude of the initial downward deflection (Fig 6F); strikingly, this progression was absent in 5-HT2B -/- mutants and the difference between the time course of the heterozygotes and homozygotes was highly significant (p≤0.0001) (Fig 6F). Similarly, the change in both the time required for ΔF/F to progress from a post-stimulus minimum to secondary maximum (Fig 6F’) and the slope of the rebound relative to epoch 1 were significantly different (Fig 6F”). Together, the differences between 5HT2B +/- and 5-HT2B -/- in the response to a dark flash (Fig 6D-D”) and these data strongly suggest that 5-HT2B mediates the effects of serotonin on the response of L2 neurons to at least one type of visual stimulus, although we cannot yet conclude whether this occurred in a cell-autonomous manner.

## Discussion

Serotonergic modulation occurs by activation of multiple receptors and associated secondary messenger cascades [56, 57]. Understanding how serotonergic signaling tunes circuit activity requires functional experiments to determine the effects on specific cells in addition to mapping the cellular and subcellular cites of individual receptors. Previous studies have mapped the expression pattern of multiple receptor subtypes to specific cells within the visual system [61, 62]. To develop the fly visual system as a new molecular-genetic model to study serotonergic neuromodulation we have confirmed the expression of the five *Drosophila* serotonin receptors in a subset of experimentally tractable cells in the lamina and used live imaging to determine their potential function. To our knowledge, these represent the first functional data for serotonergic neuromodulation of the *Drosophila* visual system with the exception of a single report in 1995 [44]. We also present the first data on the subcellular localization of serotonin receptors in the visual system. In addition, our data mapping the cellular expression pattern of the receptors are complementary to two previous transcriptomic analyses [61, 62] since we have used a different experimental approach.

We focused primarily on the lamina to identify a subset of cells that might express serotonin receptors. Our goal was to identify cells that could be used for further functional studies and the function of several lamina neurons have been previously characterized [29, 34, 82, 101]. In addition, the lamina contains a relatively small number of neurons, allowing identification based on morphology alone [71]. MiMIC-T2A-GAL4 drivers [68] are useful for replicating the endogenous expression patterns and provided a convenient method to screen for cells that express each receptor (Fig 1, S1, S3-6). To validate receptor expression before performing functional studies we used a combination of MiMIC-based reporter lines, RNA-Seq and RTq-PCR. All of these techniques indicated that 5-HT2B is expressed in L2 lamina monopolar neurons, which are involved in motion [28, 102] and contrast vision [34]. The idea that L2 cells express 5-HT2B is further supported by data from a previous report that mapped neurotransmitter receptor transcripts to cells throughout the visual system [61]. Although our RT-qPCR supports the possibility that L2 neurons might also express 5-HT7, neither we nor others detect an enrichment of 5-HT7 using RNA-Seq [61].

We observed consistent expression of 5-HT1A and 5-HT1B in T1 neurons, whose function remains poorly understood [82]. We did not detect expression of any serotonin receptors in the L1 subclass of lamina monopolar neurons, which acts in parallel to L2 for visual processing. Our data on T1 and L1 are consistent with recent transcriptional studies of these and other cells in the visual system [61, 62]. Konstantinides et al. used FACS-SMART-Seq (GSE103772) and found 5-HT1A and 5-HT1B in T1 cells, 5-HT7 in Mi1 cells, and 5-HT1A in C2 cells. Davis et al. used TAPIN-seq [61] and also reported 5-HT2B in L2 cells, 5-HT1A in C2 cells, and 5-HT7 in L5 cells, consistent with the sparse labeling experiments shown here in Fig 1 and S3. In S2 Fig we directly compare serotonin receptor expression reported for each cell type in the current study, Davis et al. [61], and Konstantinides et al. [62].

Differences in the methods used to isolate target cell populations, extract RNA, and create DNA libraries can impact sequencing results. For example, Davis et al. used TAPIN-seq to purify nuclei, subjecting tissue homogenates to two rounds of immunoprecipitation to isolate tagged nuclei before preparing RNA, a process which may lead to non-specific binding. Conversely, our use of FACS can lead to contamination by other cell types due to incomplete dissociation of GFP-tagged cells or due to non-specific labeling by the driver lines. Additionally, differences in cDNA library construction, including different approaches for poly-A mRNA enrichment, can significantly affect the results obtained in different studies. In contrast to Davis et al. we did not detect expression of serotonin receptors in photoreceptor cells or in lamina monopolar neuron L3 using MiMIC-T2A-GAL4>UAS-MCFO. These inconsistencies could be due to biological variability in levels of gene expression or infidelity in the MiMIC-based approach, which may not perfectly reflect endogenous expression due to disruption of the genetic locus. Further studies may reconcile these differences; meanwhile we suggest that using multiple, overlapping methods may be important to fully evaluate complex patterns of gene expression.

Our results suggest serotonin receptor expression in lamina glia (S4 Fig). Possible roles for glia in the visual system have been suggested previously and studies of RNA expression in glia suggest a possible enrichment in some subtypes of serotonin receptors [103, 104]. The presence of serotonin receptors in glia immediately adjacent to the basement membrane of the retina raises the possibility that they could regulate photoreceptor cells in a non cell-autonomous manner. The only prior report on acute serotonergic signaling events in the *Drosophila* visual system described the modulation of potassium channels in photoreceptors [44]. However, the ex vivo preparations used for these experiments may have included other cells or cell fragments, and unlike a previous transcriptomic study [61], we did not detect serotonin receptor expression in photoreceptors. We therefore speculate that serotonin could regulate photoreceptors non-autonomously as we report for L1 either via inputs from other neurons or perhaps glia [27, 44, 105].

Serotonergic signaling occurs via G-protein coupled receptors, which can induce immediate or long-term changes in cell physiology. We examined acute responses to serotonin receptor activation by bath applying serotonin onto optic lobe tissue. Consistent with the predicted coupling of 5-HT2B to G_q_, we found that L2 neurons respond with a robust increase in calcium measured by GCaMP6f fluorescence (Fig 4B, Fig 6A, S9A Fig) and this effect was dramatically reduced in 5-HT2B -/- flies (Fig 6B). The simplest explanation for these effects is that 5-HT2B regulates L2 in a cell autonomous manner. However, it remains possible that 5-HT2B expressed in other cells could contribute to the effects we observe. Specific knockdown of 5-HT2B in L2 neurons will be required to address this important issue, but available 5-HT2B RNAi lines have been ineffective in our hands, and additional genetic tools will be needed to perform cell type specific knock down or knock out experiments [106]. Additional experiments will also be needed to assess the potential developmental effects of the 5-HT2B mutant. Finally, we cannot completely rule out the possibility that 5-HT7 could contribute to the regulation of at least some L2 cells (see S7B Fig). These limitations aside, our data indicate that 5-HT2B plays a role in regulating one of the cells required for most visual processing in the fly.

In L1 neurons, which do not express serotonin receptors, we unexpectedly observed a large calcium response to serotonin similar to that of L2 neurons (Fig 4C). These data suggest that L1 cells are indirectly regulated by input from other cells, but further studies will be needed to determine the underlying mechanism. The most likely explanation would seem to be indirect, non-autonomous activation of L1 neurons by input from cells that express serotonin receptors either in the optic lobes or perhaps the central brain. Either synaptic or non-synaptic inputs from neighboring columnar neurons or from wide field neurons that innervate multiple layers of the medulla or glia could be responsible for these effects. TTX was included in solutions used for the experiments on L1 shown in Fig 4. TTX blocks sodium channel-based action potentials and graded potentials, which have been well documented in the insect brain including photoreceptor cells and other optic lobe neurons, and could contribute to non-autonomous regulatory effects in the presence of TTX; however, we cannot rule out the possibility that TTX affects sodium channels expressed by cells that use graded potentials [96-98]. We also note that GABAergic C2 neurons [107] express 5-HT1A (S2 Fig, S3D-E Fig) and are known to synapse onto L1 in M1 [36]. Likewise, L5 neurons express the 5-HT7 receptor (Fig 1E, S2 Fig) and reciprocally synapse with L1 neurons in M1 and M5 [36]. Alternatively, it is possible that the gap junctions that couple L1 and L2 neurons or interactions with glia could mediate indirect serotonergic regulation of L1 [28]. Regardless of the underlying mechanism, our data highlight the importance of indirect pathways for the serotonergic modulation of circuit activity.

Multiple neuroanatomical experiments, including data presented here (S5 Fig), show that serotonergic boutons are distributed within several layers of the medulla neuropil including M1, M2, M4 and the inner medulla (iM) [50, 78, 108, 109]. By contrast, they are absent from the lamina neuropil. We found that a molecularly tagged version of 5-HT2B expressed in L2 neurons was enriched in the medulla neuropil, most prominently in layer M2 (Fig 3). A tagged version of 5-HT1A expressed via the 5-HT1A locus in T1 and other cells localized to a similar site in the medulla neuropil (Fig 3). These data strongly suggest that the medulla rather than the lamina is the most relevant site for serotonergic regulation of these cells. The lamina cortex, which contains the somata of L2 and other monopolar cells is also innervated by serotonergic processes; however, to our knowledge there is no data in flies in support of the aminergic innervation of cell bodies rather than the neuropil.

Similar to most other circuits, serotonergic signaling to L2 in the medulla is likely to occur extrasynaptically. In both mammals and insects, serotonin can be released extra-synaptically through volume transmission [88, 110] or through synaptic sites [48, 111-114]. However, the majority of mammalian aminergic release sites lack an identifiable synaptic partner, and even synaptic release can lead to spillover and extrasynaptic release. Ultrastructural studies in the fly visual system have established a connectome for many neurons in the lamina and medulla including L1, L2 and T1 neurons. By contrast, the ultrastructure of serotonergic and other aminergic processes in the optic lobe of *Drosophila* are not known. Serotonergic release sites appear to lack synaptic partners in the lamina of the blowfly *Calliphora* [115] and our data using sybGRASP to test whether serotonergic neurons synapse on L2, T1, and L1 was negative (S8 Fig). It is possible that ultrastructural studies will prove more fruitful, but in the absence of true synaptic connections we anticipate that new methods may be needed to unambiguously determine the relationship of aminergic boutons to potential targets in the visual system.

In T1 neurons, we were unable to detect any acute changes in baseline calcium or voltage in response to serotonin application (Fig 4D and S9C-D Fig) and other probes (e.g., for cAMP) may be necessary to detect the acute response of T1 to serotonin. However, it is also possible that activation of 5-HT1 receptors does not induce any acute physiological response, and that more chronic indices will necessary to detect the potential effects of serotonin on T1 neurons. For now, the absence of a GCaMP6f response in T1 neurons serves as an important negative control and demonstrates the specificity of the responses observed in L1 and L2.

In *Drosophila*, L1 and L2 neurons detect changes in luminance and together are necessary for the full complement of motion vision. Both neurons receive synaptic input from photoreceptors, and respond to luminance changes with graded potentials, depolarizing in dark conditions and hyperpolarizing in light [30-32]. These two neurons feed forward into parallel pathways to enable further visual processing such as motion and contrast detection [29, 34, 101]. The modulation of visually induced calcium transients in L2 following serotonin application (Fig 5-6) suggests a role for serotonin in potentiating the response of L2-dependent visual processing pathways [31]. Our data also suggest a molecular-mechanism for previous observations made in larger insects including serotonin-induced changes field recordings in blowfly representing the output of lamina monopolar cells [53] and honeybee motion detection in the lobula [45].

The effects of serotonin on calcium signaling in L2 cells were readily apparent in experiments comparing its effects on 5-HT2B homozygous mutants versus heterozygous controls. We detected serotonin-mediated differences in both the kinetics and amplitude of L2 neurons’ response to visual stimuli and saw effects on both dark and light flash induced calcium transients in mutants (Fig 6). By contrast, experiments comparing the effects of saline versus serotonin in 5-HT2B +/+ flies showed a significant difference in only one kinetic measurement and only in response to a dark flask (Fig 5). We suggest that the 5-HT2B mutant functioned as a “sensitized genetic background” to enhance the detection of serotonin’s effects and speculate that endogenous serotonergic release and its activation of 5-HT2B may have obscured the effects of bath-applied serotonin in wild type 5-HT2B flies. A saturation point for neuromodulatory input is critical for circuit stability as has been described previously [2, 116, 117], and release of endogenous serotonin under the conditions used for these experiments might generate extracellular concentrations near the saturation point for regulating the response of L2 to visual stimuli. In the absence of a functional 5-HT2B receptor, neither endogenous nor exogenous serotonin would have any effect, thus increasing our ability to detect a difference between the 5-HT2B mutant and heterozygous controls (Fig 6). Similarly, the effects of serotonin on pathways mediated by other receptors in the visual system may only become detectable using additional receptor mutants. More generally, we suggest that our data underscore the power of a genetically sensitized background to fully understand the mechanisms underlying the aminergic regulation of circuit function.

While the precise function of serotonergic signaling in L2 neurons or elsewhere in the fly visual system remains unknown, it may allow neurons to adapt to changes in visual stimuli. We are struck by the similarities between the changes we observe during perfusion with serotonin and the graded responses of lamina neurons to stimuli of increasing contrast [31]. Changes in calcium levels at the nerve terminal suggests that serotonin might regulate inputs to L2 nerve terminals, or perhaps regulate neurotransmitter release, where increased calcium levels could drive an increase in neurotransmitter release onto postsynaptic neurons. Dissecting these potential mechanisms will require the analysis of neurons downstream of L2, but these too, may undergo direct or indirect serotonergic regulation independent of the effects of serotonin on L2 neurons [37, 118, 119]. Studies in mammals have already begun to dissect the contributions of serotonergic tuning in multiple cells within individual circuits including the visual system [6, 18, 23, 120, 121] but the way in which this information is integrated remains poorly understood. The interactions between receptors expressed on L2 and other neurons in the fly visual system provide a new framework to dissect the mechanism by which multiplexed serotonergic inputs combine to regulate circuit function.

## Conclusion

In this study we confirm that several cells in the *Drosophila* optic lobe express specific serotonin receptor subtypes. We then demonstrate that subsets of neurons involved in the initial steps of visual processing are regulated by serotonin through both presumptive cell autonomous and non-autonomous mechanisms. These data established a new platform to study the cellular mechanisms by which serotonin regulates sensory circuits.

## Supporting Information Legends

**S1 Fig. Serotonin receptors and serotonergic projections in the optic lobe**. (**A-H**) Serotonin receptor MiMIC-T2A-GAL4 lines were crossed to UAS-mCD8::RFP (visualized here in green) to identify patterns of expression in the optic lobe. (**A**) A schematic of the optic lobe neuropils including the lamina (la), medulla (me), lobula (lo) and lobula plate (lp) with the neuropil in grey and the cortex containing neuronal cell bodies in white. (**B-F**) Neuropil is labeled by anti-N-Cadherin staining (blue) to provide anatomical reference for labeled cells (green) representing 5-HT1A (**B**), 5-HT1B (**C**), 5-HT2A (**D**), 5-HT2B (**E**), and 5-HT7 (**F**). MiMIC-T2A- GAL4 line labeled projections were visible in all optic lobe neuropils including the lamina neuropil, which is enlarged in (B’), (C’), (E’) and (F’). N=4-8 and scale bars are 20 μm.

**S2 Fig. Data sets reporting evidence of serotonin receptor expression in optic lobe neurons.** The current study includes MiMIC-T2A-GAL4>MCFO for identification based on morphology (green) and FACS-SMART-Seq of L2 and T1 neurons (blue). Davis et al. 2020 employed TAPIN-Seq and reported probability of expressions (GSE116969, Table 7B) for each cell type. Serotonin receptor expression with a p>0.75 are shown (purple). Konstantinides et al. 2018 used FACS-SMART-Seq for T1, Mi1, C2 and C3 cells (GSE103772). Serotonin receptors with counts greater than 1,000 in at least two replicates are shown (orange).

**S3 Fig. Serotonin receptor MiMIC-T2A-GAL4 lines potentially label L2, C2, TMY3, and Mi1 cells.** (**A**) 5-HT2B-MiMIC-T2A-GAL4>UAS-RFP (green) was combined with ChAT-MiMIC-LexA>LexAop-GFP (magenta). Co-labeling was observed in cell bodies in the lamina cortex, shown in insets (arrowheads). (**B-C**) 5-HT7-MiMIC-T2A-GAL4 labeled cells with a morphology similar to lamina monopolar cell 5 (L5). (**D**) 5-HT1A-MiMIC-T2A-GAL4>MCFO labeled C2-like cells in the lamina neuropil. (**E**) Colocalization was observed between 5-HT1A MiMIC-T2A-GAL4 and GAD1 MiMIC-T2A-LexA in the lamina neuropil and cell bodies adjacent to the lobula plate (arrowhead, E). (**F-G**) 5-HT7-MiMIC-GAL4>MCFO (F) and 5-HT1A-MiMIC-T2A-GAL4>MCFO (G) labeled cells with morphology similar to TmY3. (**H**) 5-HT7-MiMIC-GAL4>MCFO also labeled cells that resembled Mi1. For co-labeling in (A) and (E), N=4-8 brains per condition. For MCFO, L5-like cells in (B-C) were observed in 7/13 brains, C2 cells in (D) were observed in 3/31 brains, TMY3 cells in (F) were observed in 4/13 brains, TmY3 cells in (G) were observed in 5/31 brains and Mi1 cells in (H) were observed in 6/13 brains. Scale bars are 20 µm for (A, D-H) and 10 µm for (B-C).

**S4 Fig. 5-H T2A labeling in lamina cortex may represent glia cells.** (**A**) 5-HT2A-GAL4>MCFO epitopes V5 (green) and HA (magenta) label unidentified cells confined the distal lamina cortex. (**B**) 5-HT2A-T2A-GAL4>UAS-mCD8::GFP (green) labels cells in the lamina cortex in close proximity to nuclei labeled with repo antibody (magenta). Neuropil is labeled by anti-N-Cadherin staining (blue) to provide anatomical reference. N=18 brains for (A) and N=4 brains from (B). Scale bars are 20 um.

**S5 Fig. Serotonin receptor 5-HT1B co-labels with serotonin immunoreactive sites in optic lobe and cell bodies in the central brains.** Anti-serotonin immunolabeling (magenta) was used to identify serotonergic cells and projections in (A-F). (**A**) Serotonin immunoreactive sites (magenta) were visible in the optic lobe neuropils: lamina (la), medulla (me), lobula (lo) and lobula plate (lp). (**A’-A” and C-C”**) 5-HT1B-MiMIC-T2A-GAL4>UAS-mCD8::GFP labeled cells throughout the optic lobe with close apposition to serotonergic boutons. A neuron in serotonergic cell cluster LP2 is also labeled by 5-HT1B driven GFP (arrowhead, A-A”). (**B**) A schematic of the optic lobe and its major neuropils. (**E-E”**) Serotonin receptor MiMIC-T2A-GAL4 lines were crossed to UAS-MCFO-1 to label individual cells. Using 5-HT1B-MiMIC-T2A-GAL4>MCFO (green), we observed co-labeling between MCFO-labeled cells and serotonergic boutons (magenta) processes in the inner medulla (iM), medulla layer 4 (M4), and lobula (lo). (**D**) A schematic of the fly brain with dashed lines showing the approximate anatomical locations for (C) and (E). (**F-F”**) Anti-serotonin immunolabeling (magenta) co-labeled with 5-HT1B-MiMIC-T2A-GAL4>UAS-mCD8::GFP labeled cell bodies in the central brain. 5-HT1B-labeled kenyon cells (KC) are labeled for anatomical reference in (F”). (**G**) The approximate anatomical location for images in (F-F”) are shown in the boundaries of the dashed line. Serotonin co-labeling was performed N=5 for 5-HT1B>GFP (A-A”, C-C”, and F-F”) and N=6 brains for 5-HT1B>MCFO (E-E”). Scale bars are 20 μm.

**S6 Fig. Serotonin receptor 5-HT1A co-labels with serotonin immunoreactive sites in optic lobe and cell bodies in the central brain.** (**A**) Schematic of the optic lobe neuropils—lamina (la), medulla (me), lobula (lo) and lobula plate (lp)—and serotonergic PLP cells. (**B-B”**) 5-HT1A-MiMIC-T2A-GAL4 driving UAS-mCD8::GFP (green) was co-stained with anti-serotonin immunolabeling (magenta) to map potential autoreceptors to specific cell clusters in the central brain. PLP neurons co-labeling for 5-HT1A labeling and anti-serotonin immunolabeling are indicated by the arrowhead. (**C-E**) Anterior to posterior images taken in the same brain show several serotonergic cell clusters expressing 5-HT1A (labeled at arrowheads). 5-HT1A-labeled mushroom body (MB) and kenyon cells (KC) are labeled for anatomical reference in (**C** and **E**). (**F**) A cartoon of the fly brain with dashed lines to indicate the approximate anatomical location for (C-E). Scale bars are 20 μm and N=6.

**S7 Fig. RT-qPCR shows L2 neurons express 5-HT2B and T1 neurons express both 5-HT1A and 5-HT1B serotonin receptors.** Enrichment (i.e., fold change) was calculated for cDNA from GFP-labeled cell isolates relative to pooled, unlabeled optic lobe cell isolates using the comparative CT method. (**A**) RT-qPCR performed on cDNA from isolated T1 neurons expressing GFP showed enrichment for serotonin receptors 5-HT1A and 5-HT1B relative to other GFP-negative cells from the optic lobe. (**B**) FACS isolates from L2 cells showed enrichment of 5-HT2B and 5-HT7 in RT-qPCR. (**C**) L1 RT-qPCR enrichment was not detectable for any serotonin receptors. RT-qPCR error bars represent mean±SEM. N=3-6 biological replicates pooled from 18-40 brains per replicate.

**S8 Fig**. **Serotonergic neurons do not show sybGRASP signal with postsynaptic T1, L2 or L1 neurons in the medulla.** SybGRASP was used to probe whether serotonergic neurons make synaptic contacts onto L2, T1 or L1 neurons. L2 neurons are known to synapse onto T1 projections in the medulla and we used this connection as a positive control. (**A-A’**) SybGRASP was observed with L2 split-GAL4 presynaptic to T1-LexA in M2. The dashed inset in (**A**) is shown in (**A’**). (**B-E**) A SerT-GAL4 driver was used to express the pre-synaptic portion of GFP in serotonergic neurons and LexA drivers were used to express the postsynaptic portion of GFP in L2 (**B-C**) L1 (**D**) or T1 (**E**) as indicated. No sybGRASP signal was detected in the medulla when SerT was presynaptic to L2 (**B**) however, occasional sparse GFP puncta (arrowhead) were visible in the lamina (**C**). When SerT-GAL4 neurons were presynaptic to L1 (**D**) or T1 (**E**) neurons, we did not detect a sybGRASP signal in either the lamina or medulla. All tissue was labeled with primary antibodies to both 5-HT (magenta) and GFP (green). N=7-10 brains. Scale bars are 15 µm (**A**, **B**, **D-E**); 5 µm (**A”** and **C-C”**).

**S9 Fig. Individual traces for serotonin bath application experiments.** (**A-D**) Individual traces representing all experiments (Fig 4) for serotonin or saline controls with L2, L1 or T1 split-GAL4>GCaMP6f or T1 split-GAL4>ArcLight. For all experiments, the first 60 s of baseline is not shown; traces represent data recorded following a switch to saline with serotonin or saline alone. The length of time for the switch to complete was estimated to be 105 s, but the first 60 s of the recording are not shown so the switch occurs at 45 s on the x axis in A-D. Saline controls are gray, serotonin exposed preps are colored, and the dark line represents the mean (**A**) L2>GCaMP6f experiments, along with (**B**) L1>GCaMP6f, show an increase in calcium following serotonin application as compared to saline controls (L2, p=00095; L1, p=0.02). (**C, D**) T1 cells show no significant change with either GCaMP6f (**C**) or ArcLight (**D**) relative to saline (p>0.05) For bath application experiments (**A-D**), N=4-8 individual flies.

**S1 Table.** RNA-Seq Serotonin Receptor TPMs, averages and standard deviations.

**S2 Table.** RT-qPCR Threshold Cycle (CT) measurements and calculated enrichment for FACS-isolated T1, L2, and L1 samples as shown in S7 Fig. Enrichment (i.e., fold change) was calculated for cDNA from GFP-labeled cell isolates relative to pooled, unlabeled optic lobe cell isolates using the comparative CT method.

**S3 Table.** Fly strains used in this study.

**S4 Table.** RT-qPCR primer sequences and mRNA (cDNA) target information.

## Methods

### Fly Husbandry and Genetic Lines

Flies were maintained on a standard cornmeal and molasses-based agar media with a 12:12 hour light/dark cycle at room temperature (22-25**°**C). All fly strains used in this study are listed in S3 Table. Serotonin receptor MiMIC-T2A-GAL4 lines described in [68] were a gift from Herman Dierick (Baylor College of Medicine), and include 5-HT1A-T2A-GAL4^MI01468^, 5-HT1A-T2A-GAL4^MI01140^, 5-HT1A-T2A-GAL4^MI04464^, 5-HT1B-T2A-GAL4^MI05213^, 5-HT2A-T2A-GAL4 ^MI0459^, 5-HT2A-GAL4^MI03299^, 5-HT2B-T2A-GAL4^MI06500^, 5-HT2B-T2A-GAL4^MI5208^, 5-HT2B-GAL4^MI7403^, and 5-HT7-GAL4 ^MI00215^. L1 and T1 split-GAL4 lines [82], as well as unpublished LexA lines for L1 and T1, were provided by Aljoscha Nern (HHMI/Janelia Research Campus). S.L. Zipursky (UCLA) generously provided L2-split-GAL4 and L2-LexA (RRID:BDSC_52510). Yi Rao (Peking University) generously shared 5-HT2B-KO-GAL4 SII (5-HT2B mutant) [85], 5-HT1A::sfGFP [84], and UAS-5-HT2B::sfGFP [85].

SerT-GAL4 (RRID:BDSC_38764), GAD1 Trojan LexA (RRID:BDSC_60324) and ChAT Trojan LexA (RRID:BDSC_60319) were obtained from Bloomington *Drosophila* Stock Center (BDSC) at Indiana University (Bloomington, IN, USA). Additional reporters from BDSC include: UAS-mCD8::GFP (RRID:BDSC_5137), UAS-MCFO-1 (RRID:BDSC_64085), UAS-GCaMP6f (RRID:BDSC_42747), UAS-ArcLight (RRID:BDSC_51056), UAS-mCD8::RFP, LexAop-mCD8::GFP (RRID:BDSC_32229), and UAS-nSyb::GFP1-10, LexAop-CD4:GFP11 (RRID:BDSC_64314; provided through S.L. Zipursky (UCLA)).

**S3 Table.** Animal strains used in this study.

### Immunofluorescent Labeling and Imaging

Flies were dissected 5-10 days after eclosion, and equal numbers of males and females were used for all experiments unless otherwise noted. Brains were dissected in ice-cold PBS (Alfa Aesar, Cat#J62036, Tewksbury, MA), then fixed in 4% paraformaldehyde (FisherScientific, Cat#50-980-493, Waltham, MA) in PBS with 0.3% Triton X-100 (Millipore Sigma, Cat#X100, Burlington, MA) (PBST) for one hour at room temperature. After fixation, brains were washed three times with PBST for 10 minutes, then blocked for 30 minutes in PBST containing 0.5% normal goat serum (NGS) (Cayman Chemical, Cat#10006577, Ann Arbor, MA). Antibodies were diluted in 0.5% NGS/PBST. Primary antibodies were incubated with the tissue overnight at 4**°**C. The next day, the brains were washed three times with PBST for 10 minutes, then incubated with secondary antibodies for 2 hours in the dark at room temperature. Brains were washed three times with PBST for 10 minutes before mounting.

For frontal mounting, brains were washed with 60% and 80% glycerol (Millipore Sigma, Cat#G5516) and mounted in Fluoromount-G (SouthernBiotech, Cat#0100-01, Birmingham, AL). For dorsal-ventral mounting, brains were fixed in 2% PFA/PBST overnight. The next day, brains were washed three times with PBST. Brains were dehydrated with a series of 10 min ethanol baths of increasing concentrations (30%, 50%, 75%, 95%, 100%, 100%, and 100%). Brains were then transferred to 100% xylene before mounting in DPX (FisherScientific, Cat#50-980-370).

Serotonin immunolabeling was performed with 1:25 rat anti-serotonin (Millipore Sigma, Cat#MAB352, RRID:AB_11213564), 1:1000 rabbit anti-serotonin (ImmunoStar, Cat#20080, Hudson, WI, RRID:AB_572263) or 1:1000 goat anti-serotonin (ImmunoStar, Cat#20079, RRID:AB_572262). Where noted, GFP was labeled with 1:250 mouse anti-GFP (Sigma-Aldrich, Cat#G6539, RRID:AB_259941; or, ThermoFisher, Waltham, MA, Cat#A-11120, RRID:AB_221568). Secondary antibodies were used at 1:400 and include: Donkey anti-mouse Alexa Fluor 488, Donkey anti-Rabbit Alexa Fluor 594 or Alexa Fluor Donkey anti-rat 647 (Jackson ImmunoResearch Laboratories, Westgrove, PA, Cat#715-545-151, # 711-585-152, # 712-605-153) or Alexa Fluor 555 (Life Technologies, ThermoFisher, Cat#A-21428).

When serotonin receptor MiMIC-GAL4 lines were combined with ChAT or GAD1 MiMIC-LexA (S3 Fig), brains were processed and imaged as described in Sizemore and Dacks 2016 [122]. MultiColor FlpOut (MCFO-1) sparse labeling was induced by heat activation at 37**°**C for 10-15 minutes at least 2 days prior to dissection as described [73]. Primary antibodies included 1:300 rabbit anti-HA (Cell Signaling Technology, Cat#3724, Danvers, MA, RRID:AB_1549585), 1:150 rat anti-FLAG (Novus, Littleton, CA, Cat#NBP1-06712, RRID:AB_1625982), and 1:400 mouse anti-V5::Dylight-550 (Bio-Rad, Hercules, CA, Cat#MCA1360D550GA, RRID:AB_2687576). The rat anti-N-Cadherin (DN-Ex#8) and mouse anti-repo (8D12 concentrate) was obtained from the Developmental Studies Hybridoma Bank, created by the NICHD of the NIH and maintained at The University of Iowa, Department of Biology, Iowa City, IA 52242. Secondary antibodies used for MCFO are listed above. N-Synaptobrevin GFP Reconstitution Across Synaptic Partners (sybGRASP) flies [86] were dissected, fixed and immunolabeled as described above, without KCl induction. The tissue was labeled with mouse antiserum specific to reconstituted GFP (1:250; Sigma-Aldrich, Cat#G6539, RRID:AB_259941) [123] and either anti-serotonin (antibodies listed above).

Imaging was performed with a Zeiss LSM 880 Confocal with Airyscan (Zeiss, Oberkochen, Germany) using a 40x water or 63x oil immersion objective. Post-hoc processing of images was done with Fiji [125] or Adobe Photoshop (Adobe, San Jose, CA).

### FACs and RNA Extraction

L2 and L1 neurons were labeled using split-GAL4 drivers combined with UAS-mCD8::GFP (RRID:BDSC_5137). For RNA-Seq in Fig 2, N=3 T1-LexA samples were tested. For RT-qPCR in S7 Fig, we included N=3 T1-split-GLA4 samples and N=3 T1-LexA samples. Brains were dissected on the day of eclosion and optic lobes were dissociated according to previously published methods [126]. The dissociated optic lobe cells were separated by fluorescence-activated cell sorting (FACS) into GFP-positive and GFP-negative isolates using a BD FACS Aria II high-speed cell sorter in collaboration with the UCLA Jonsson Comprehensive Cancer Center (JCCC) and Center for AIDS Research Flow Cytometry Core Facility (http://cyto.mednet.ucla.edu/home.html). For FACS, each experiment was performed with 18-40 brains, and yielded between 1,700-7,800 GFP^+^ cells. RNA was extracted from isolated cells with ARCTURUS® PicoPure® RNA Isolation Kit (ThermoFisher, KIT0204) or RNeasy Plus Micro Kit (QIAGEN, 74034).

### RT-qPCR

RNA extracted from FACS isolates was reverse transcribed using SuperScript III (Invitrogen, ThermoFisher, Cat#18080093). RT-qPCR was performed for receptor cDNA using validated primers (S4 Table) and SYBR Green Power PCR Mix (Applied Biosystems, ThermoFisher) on an iQ5 real-time qPCR detection system (Bio-Rad). Primers were designed using Primer-Blast (https://www.ncbi.nlm.nih.gov/tools/primer-blast/) or were from the DGRC FlyPrimerBank [127]; oligonucleotides were obtained from Integrated DNA Technologies (Coralville, Iowa). Primer pairs were validated to amplify a single product, verified by a single melting temperature and single band on an electrophoresis gel. The efficiency for each primer pair was between 85-115%. Comparisons between GFP^+^ and GFP^-^ samples were calculated as enrichment (i.e., fold change) using the comparative CT method [83]. A zero value was imputed for samples with no amplification (i.e., no CT value). Raw CT values are shown in S2 Table.

### RNA-Seq

RNA-Seq was performed using a SMART-Seq protocol adapted from [126, 128, 129]. Libraries were constructed using the SMART-seq v4 Ultra Low-input RNA sequencing kit with Nextera XT (Takara Bio). Paired-end sequencing was conducted by the UCLA genomic core facility (https://www.semel.ucla.edu/ungc/services). After demultiplexing, we obtained between 39-270 (average 105) million reads per sample. Quality control was performed on base qualities and nucleotide composition of sequences. Alignment to the *Drosophila melanogaster* genome (BDGP6) was performed using the STAR spliced read aligner [130] with default parameters. Additional QC was performed after the alignment to examine the following: level of mismatch rate, mapping rate to the whole genome, repeats, chromosomes, and key transcriptomic regions (exons, introns, UTRs, genes). Between 75-85% of the reads mapped uniquely to the fly genome. Total counts of read fragments aligned to candidate gene regions within the reference gene annotation were derived using HTSeq program and used as a basis for the quantification of gene expression. Only uniquely mapped reads were used for subsequent analyses. Following alignment and read quantification, we performed quality control using a variety of indices, including consistency of replicates and average gene coverage. For Fig 2, L2 samples were run in two separate sequencing runs and we did not perform corrections for any potential batch effects. Data is shown as Transcripts Per Million (TPMs).

### Live Cell Imaging

Calcium imaging was performed as previously described [131]. Briefly, flies were anesthetized at 4°C and placed into a chemically etched metal shim within a larger custom-built fly holder. The fly holder was based on a previously described design [132]. The head capsule and the thorax were glued to the metal shim using UV-curable glue (www.esslinger.com). The legs, proboscis and antennae were immobilized using beeswax applied with a heated metal probe (Waxelectric-1, Renfert). The head capsule was immersed in insect saline (103 mM NaCl, 3 mM KCl, 1.5mM CaCl2, 4 mM MgCl2, 26 mM NaHCO3, 1 mM NaH2PO4, 10 mM trehalose, 10 mM glucose, 5 mM TES, 2 mM sucrose) [133]. A small window on the right rear head capsule was opened using sharp forceps (Dumont, #5SF). Muscles and fat covering the optic lobe were cleared before placing the fly under the 2-photon microscope (VIVO, 3i: Intelligent Imaging Innovations, Denver, CO). Neurons expressing GCaMP6f were imaged at 920-nm using a Ti:Sapphire Laser (Chameleon Vision, Coherent). Images were acquired at 10-20 frames/s for Fig 4 and 25-30 frames/s for Fig 5-6 live imaging. Only female flies were used for live imaging experiments.

A custom-built gravity perfusion system was used for bath application of either serotonin or saline control to the fly’s exposed optic lobe for Fig 4. For Fig 4 and S9 Fig, the tissue was first perfused with insect saline containing 1μm tetrodotoxin citrate (TTX) (Alomone Labs, Jerusalem, Israel, Cat#T-550) for at least 5 minutes at 2 mL/min, prior to each recording. TTX remained present throughout the experiment. To examine the effects of serotonin on calcium levels, baseline GCaMP6f fluorescence was recorded for one minute before switching to the second input containing either 100µM serotonin hydrochloride (Sigma Aldrich, Cat# H9523) or saline alone for an additional five minutes of recording. Due to perfusion tubing length and dead volume, the perfusion switch took approximately 105 s to reach the tissue. Fig 4 does not show the first minute of the recording, so the solution switch occurs at approximately 45 s on the x axis. For more precise control of perfusion solutions, we used a programmable valve controller (VC-6, Warner Instruments, Hamden, CT) for Fig 5-6 visual experiments (see details below).

### Visual Stimulus Experiments

Visual stimuli were shown using an arena composed of 48 eight by eight-pixel LED panels, at 470 nm (Adafruit, NY, NY). The panels were assembled into a curved display that extends 216° along the azimuth and ±35° in elevation. Each pixel subtended an angle of 2.2° on the retina at the equatorial axis. To prevent spurious excitation of the imaging photomultiplier tubes, three layers of blue filter (Rosco no. 59 Indigo) were placed over the LED display.

Each stimulus consisted of a brief increment (light flash) or decrement (dark flash) of the entire display for 100 ms, before returning to a mid-intensity brightness for 4.9 s. Images were acquired at 25-30 frames/s for Fig 5-6 visual stimulation experiments. Stimuli were presented in sets of six bright and six dark flashes randomly shuffled for each minute of the experiment. Responses were then pooled for each minute. During the first minute (“epoch 1” in Figs 5-6), and prior to imaging, the tissue was perfused with saline for a baseline recording. At the end of the first minute, a valve controller (VC-6, Warner Instruments, Hamden, CT) activated by a TTL signal switched the perfusion to either saline with 100 µM serotonin or saline alone; imaging then continued for an additional five minutes, for a total of one baseline set and five post-switch sets of stimuli. The perfusion switch took approximately 45 s to reach the tissue using the programmable valve system.

### Analysis

Calcium imaging data were analyzed with Matlab R2017a (Mathworks, Natick, MA). Post hoc, recordings were corrected for movement of the brain within the imaging plane using a custom algorithm [134]. Regions of interest (ROIs) were found semi-automatically for data in Figs 4-6: first, the median intensity of all pixels across all image frames was found; this value was used as a threshold and all pixels with mean intensity below the threshold, typically within the image background, were discarded. The 1-D time series of intensity for each remaining pixel was then extracted. K-means clustering was used to identify pixels with similar activity over the course of the experiment: three clusters were identified and the cluster with the highest number of pixels was retained. This reliably identified the pixels within active neurons in the imaging data and aided in identifying preparations with out-of-plane movement, which were discarded.

For baseline activity experiments (Fig 4), the remaining cluster was used as a single ROI and the mean intensity within the ROI was found for each image frame to produce a single time-series for the entire experiment. For visual response experiments (Figs 5-6), pixels within the remaining cluster were automatically divided into groups corresponding to individual L2 terminals using a watershed transform. The mean intensity within each ROI was found for each image frame to produce a single time-series for the entire experiment, and the time-series for all terminal ROIs within an individual animal were then averaged. For Fig 6, ROIs of L2 terminals were first identified automatically, as above, then manually selected individually according to layer position because the 5-HT2B GAL4 SII mutant line labeled other cells in addition to L2 neurons.

Approximately half of the bath application recordings showed oscillations in activity due to slow, periodic movement of the brain at around 0.04 Hz; we applied a second-order notch filter at this frequency with a bandwidth of 0.005 Hz to remove these oscillations. For the bath application experiments (Fig 4), we plotted ΔF/F, defined as (F_t_-F_0_)/F_0_, where F_t_ is the mean fluorescence in the ROI at the indicated time and F_0_ is the mean value of F_t_ during 60 seconds of baseline activity at the beginning of the experiment and prior to the change in perfusion. For the visual stimulus experiments (Figs 5-6), we again plotted ΔF/F, defined as (F_t_-F_0_)/F_0_, where F_t_ is the mean fluorescence across all individual terminal ROIs at the indicated time and F_0_ is the mean of 30 seconds of non-consecutive baseline activity between stimulus presentations during epoch 1 at the beginning of the experiment and prior to the change in perfusion (Fig 5A). For the stimulus response plots (Figs 5B, D, 6C, E), we found the average ΔF/F time-series within each epoch for each fly after subtracting the average pre-stimulus baseline activity level (0.5 s preceding each flash stimulus) from each time-series, so that all responses started aligned at 0 ΔF/F. For further analysis (Figs 5C, E, 6D, F), we found the changes in response amplitude across epochs, defined for the dark stimulus presentation (top) as the difference between the pre-stimulus baseline and the maximum ΔF/F value that occurred up to 1.75 s after cessation of the 0.1 s flash, and for the light stimulus (bottom) as the minimum ΔF/F value that occurred in the same period of time. For each epoch, we subtracted the value of the responses during epoch 1 at the beginning of the experiment and prior to the change in perfusion, in order to find the change in amplitude relative to epoch 1. We followed a similar procedure for the analysis of peak-peak time (Figs 5C’, E’, 6D’, F’), defined as the length of time between the minimum and maximum ΔF/F values that occurred within 1.75 s after cessation of the 0.1 s flash. The slope of the transient decay was also calculated (Figs 5C”, E”, 6D”, F”).

To examine the baseline changes in fluorescence in visual experiments shown in Figs 6A-B responses to the visual stimuli were removed using a series of second-order notch filters at 0.19-0.21 Hz and 0.38-0.42 Hz.

### Statistical Tests

For Fig 5 (C-C”, E-E”) and Fig 6 (D-D”, E-E”) comparisons are two-way repeated measure ANOVA (brackets show interactions between time and genotype) and Sidak’s multiple comparisons tests, p≤0.05 *, p≤0.01**, p≤0.001***, p≤0.0001****. These tests were performed using Graphpad Prism Software (San Diego, CA). Differences in baseline calcium shown in Fig 6A-B were calculated by two-tailed Wilcoxon rank sum tests in Matlab R2017a.

### Replicates

Each biological replicate (N) represents one fly, except for RT-qPCR and RNA-Seq (Fig 2, and S7 Fig E) where each biological replicate was pooled from 18+ flies. RT-qPCR experiments include 3 technical replicates, which are averaged to represent a single biological replicate. Animals from at least 3 crosses were used for each experiment. Data for each experiment was collected over 2-6 months in at least 3 experiments. No outliers were removed from any data set. Live imaging recordings with too much movement were excluded and not analyzed.

## Acknowledgements

We thank members of Larry Zipursky’s lab (UCLA) including Elizabeth Zuniga Sanchez, Liming Tan, and Juyoun Yoo for advice on FACS, RNA extraction, RT-qPCR and SMART-Seq. Thank you to Deanne Umbay (Brown University) and Nikoo Dalili (UCLA) for their assistance. We thank members of the Frye and Krantz labs for helpful discussions, and Aljoscha Nern (HHMI/Janelia Research Campus) and Herman Dierick (Baylor) for generously supplying fly lines.

