## Supplemental Information for "Serotonergic modulation of visual neurons in *Drosophila melanogaster*"

**S1 Fig. Serotonin receptors and serotonergic projections in the optic lobe**. (**A-H**) Serotonin receptor MiMIC-T2A-GAL4 lines were crossed to UAS-mCD8::RFP (visualized here in green) to identify patterns of expression in the optic lobe. (**A**) A schematic of the optic lobe neuropils including the lamina (la), medulla (me), lobula (lo) and lobula plate (lp) with the neuropil in grey and the cortex containing neuronal cell bodies in white. (**B-F**) Neuropil is labeled by anti-N-Cadherin staining (blue) to provide anatomical reference for labeled cells (green) representing 5-HT1A (**B**), 5-HT1B (**C**), 5-HT2A (**D**), 5-HT2B (**E**), and 5-HT7 (**F**). MiMIC-T2A-GAL4 line labeled projections were visible in all optic lobe neuropils including the lamina neuropil, which is enlarged in (B'), (C'), (E') and (F'). N=4-8 and scale bars are 20 μm.

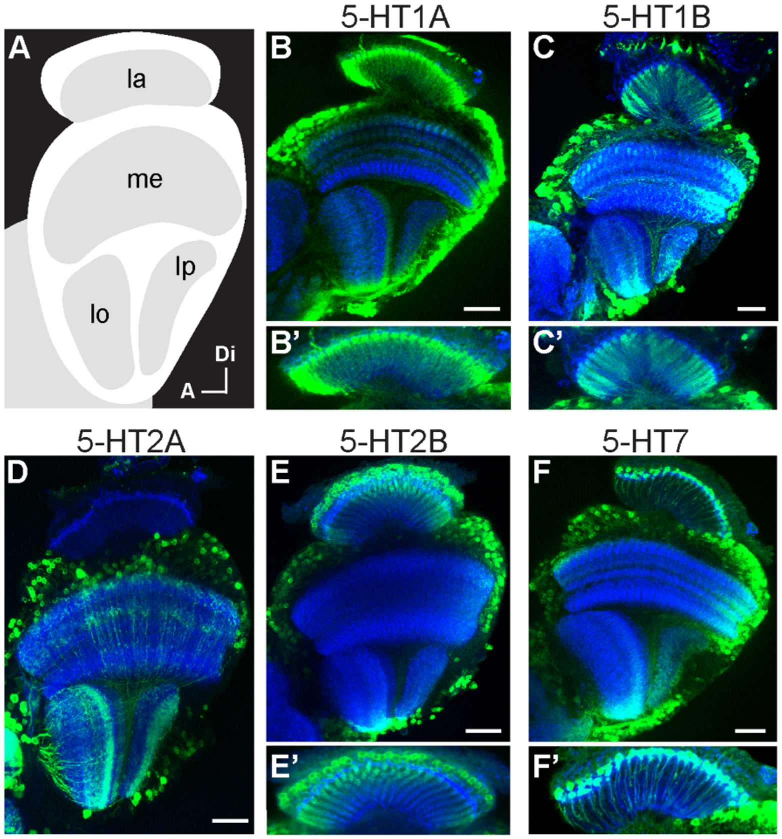

**S2 Fig. Data sets reporting evidence of serotonin receptor expression in optic lobe neurons.** The current study includes MiMIC-T2A-GAL4>MCFO for identification based on morphology (green) and FACS-SMART-Seq of L2 and T1 neurons (blue). Davis et al. 2020 employed TAPIN-Seq and reported probability of expressions (GSE116969, Table 7B) for each cell type. Serotonin receptor expression with a p>0.75 are shown (purple). Konstantinides et al. 2018 used FACS-SMART-Seq for T1, Mi1, C2 and C3 cells (GSE103772). Serotonin receptors with counts greater than 1,000 in at least two replicates are shown (orange).

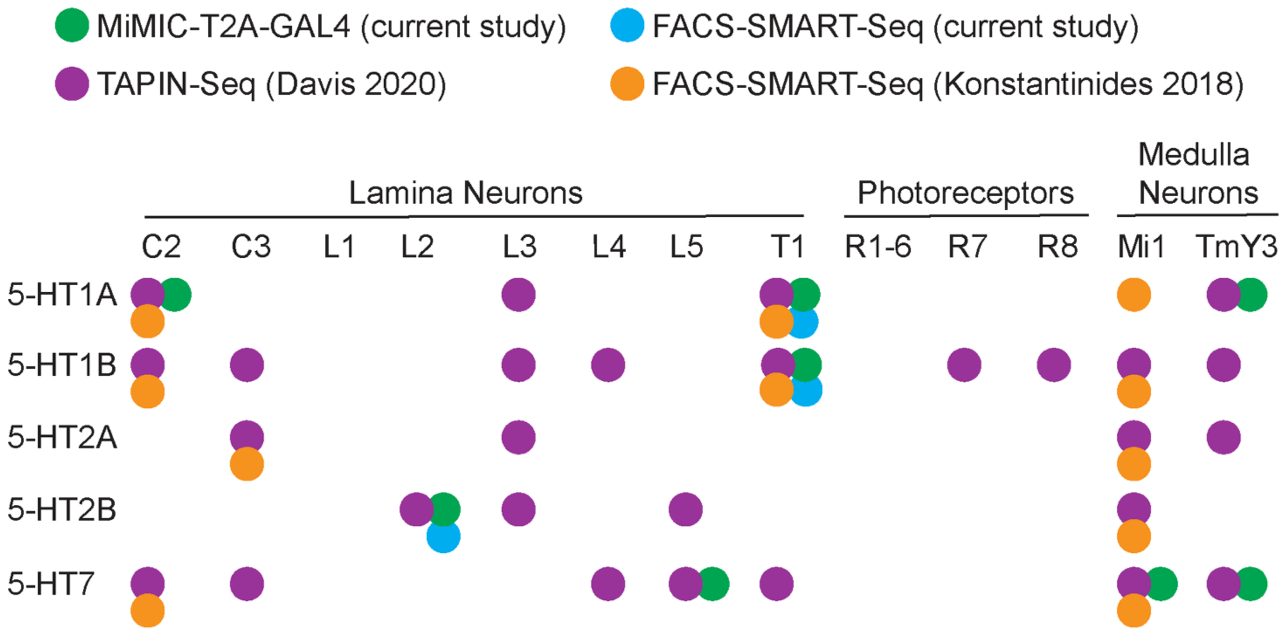

**S3 Fig. Serotonin receptor MiMIC-T2A-GAL4 lines potentially label L2, C2, TMY3, and Mi1 cells.** (**A**) 5-HT2B-MiMIC-T2A-GAL4>UAS-RFP (green) was combined with ChAT-MiMIC-LexA>LexAop-GFP (magenta). Co-labeling was observed in cell bodies in the lamina cortex, shown in insets (arrowheads). (**B-C**) 5-HT7-MiMIC-T2A-GAL4 labeled cells with a morphology similar to lamina monopolar cell 5 (L5). (**D**) 5-HT1A-MiMIC-T2A-GAL4>MCFO labeled C2-like cells in the lamina neuropil. (**E**) Colocalization was observed between 5-HT1A MiMIC-T2A-GAL4 and GAD1 MiMIC-T2A-LexA in the lamina neuropil and cell bodies adjacent to the lobula plate (arrowhead, E). (**F-G**) 5-HT7-MiMIC-GAL4>MCFO (F) and 5-HT1A-MiMIC-T2A-GAL4>MCFO (G) labeled cells with morphology similar to TmY3. (**H**) 5-HT7-MiMIC-GAL4>MCFO also labeled cells that resembled Mi1. For co-labeling in (A) and (E), N=4-8 brains per condition. For MCFO, L5-like cells in (B-C) were observed in 7/13 brains, C2 cells in (D) were observed in 3/31 brains, TMY3 cells in (F) were observed in 4/13 brains, TmY3 cells in (G) were observed in 5/31 brains and Mi1 cells in (H) were observed in 6/13 brains. Scale bars are 20 μm for (A, D-H) and 10 μm for (B-C).

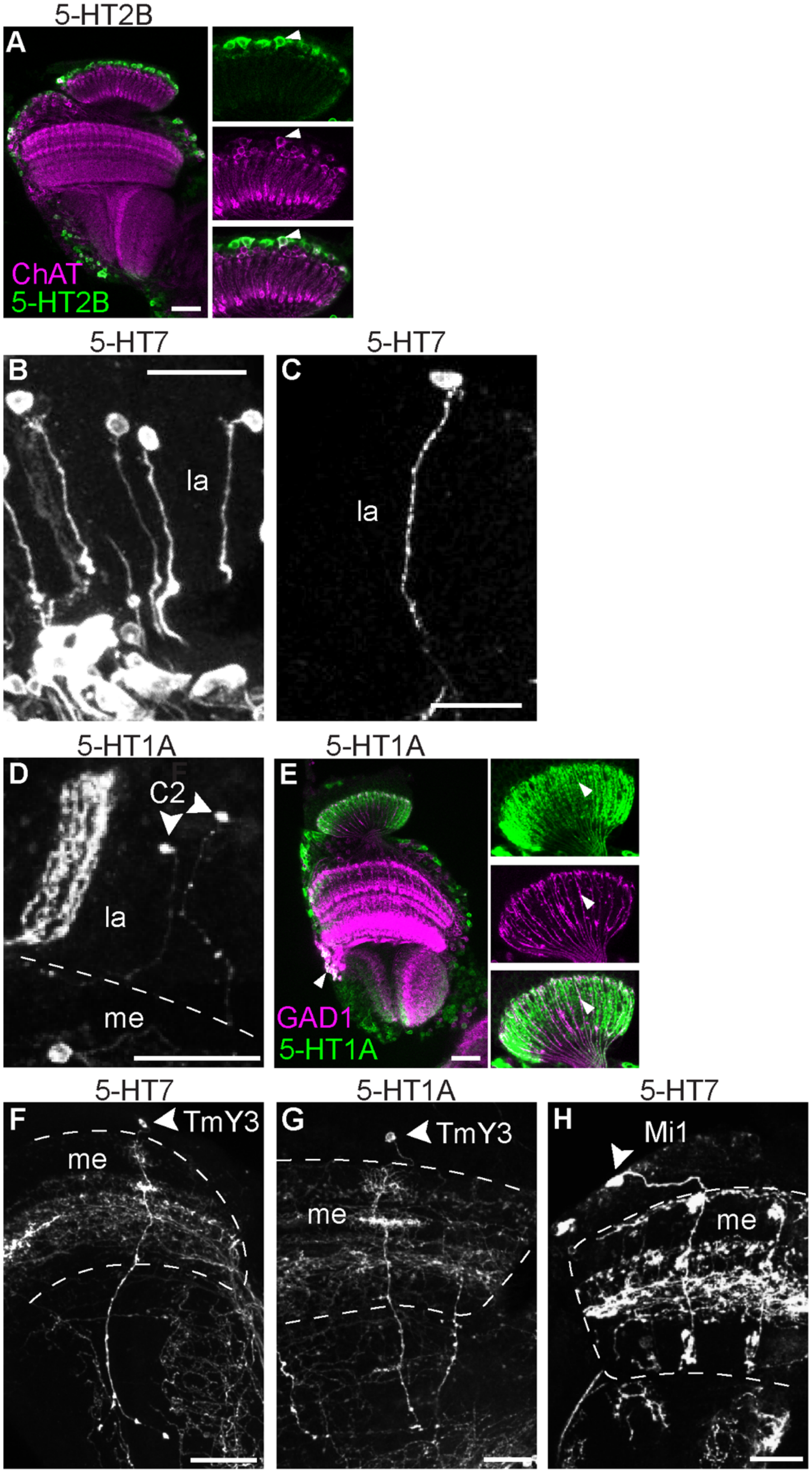
5-HT2B
A
ChAT
5-HT2B
5-HT7
B
la
5-HT7
C
la
5-HT1A
D
C2
la
me
5-HT1A
E
GAD1
5-HT1A
5-HT7
F
TmY3
me
5-HT1A
G
TmY3
me
5-HT7
H
Mi1
me

**S4 Fig. 5-HT2A labeling in lamina cortex may represent glia cells.** (**A**) 5-HT2A-GAL4>MCFO epitopes V5 (green) and HA (magenta) label unidentified cells confined the distal lamina cortex. (**B**) 5-HT2A-T2A-GAL4>UAS-mCD8::GFP (green) labels cells in the lamina cortex in close proximity to nuclei labeled with repo antibody (magenta). Neuropil is labeled by anti-N-Cadherin staining (blue) to provide anatomical reference. N=18 brains for (A) and N=4 brains from (B). Scale bars are 20 um.

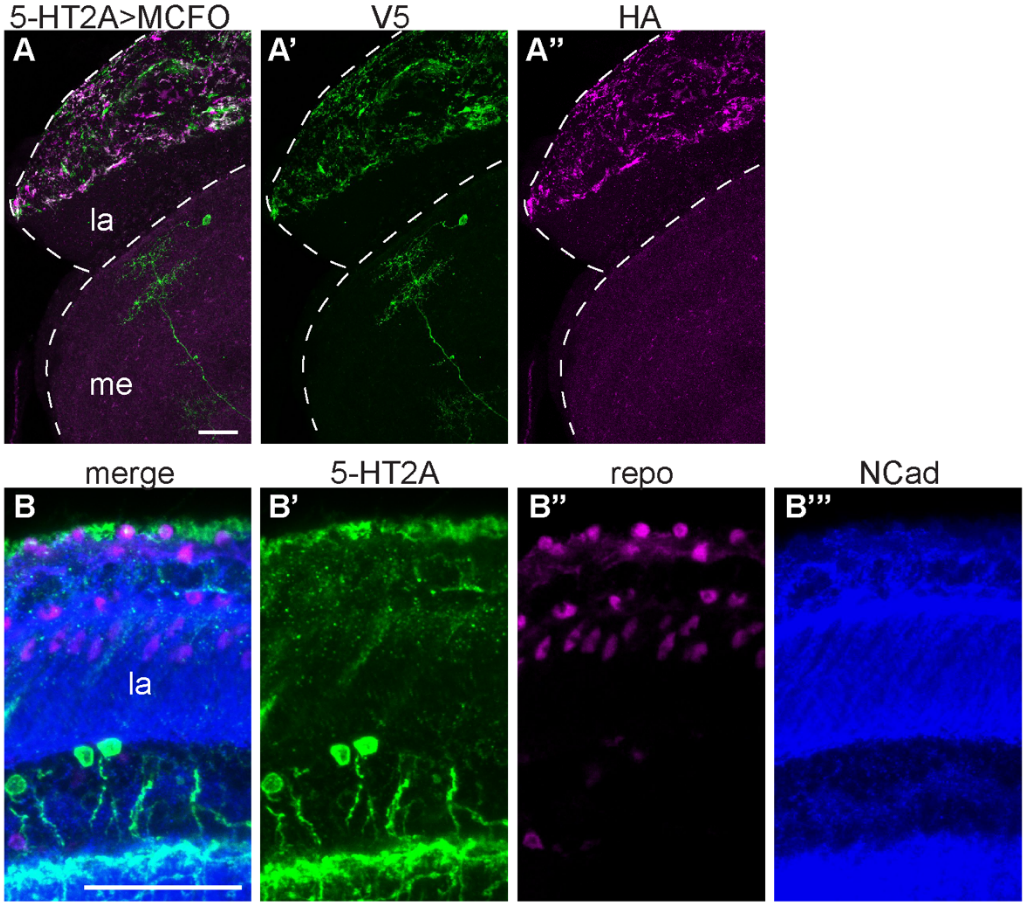

**S5 Fig. Serotonin receptor 5-HT1B co-labels with serotonin immunoreactive sites in optic lobe and cell bodies in the central brains.** Anti-serotonin immunolabeling (magenta) was used to identify serotonergic cells and projections in (A-F). (**A**) Serotonin immunoreactive sites (magenta) were visible in the optic lobe neuropils: lamina (la), medulla (me), lobula (lo) and lobula plate (lp). (**A'-A" and C-C"**) 5-HT1B-MiMIC-T2A-GAL4>UAS-mCD8::GFP labeled cells throughout the optic lobe with close apposition to serotonergic boutons. A neuron in serotonergic cell cluster LP2 is also labeled by 5-HT1B driven GFP (arrowhead, A-A"). (**B**) A schematic of the optic lobe and its major neuropils. (**E-E"**) Serotonin receptor MiMIC-T2A-GAL4 lines were crossed to UAS-MCFO-1 to label individual cells. Using 5-HT1B-MiMIC-T2A-GAL4>MCFO (green), we observed co-labeling between MCFO-labeled cells and serotonergic boutons (magenta) processes in the inner medulla (iM), medulla layer 4 (M4), and lobula (lo). (**D**) A schematic of the fly brain with dashed lines showing the approximate anatomical locations for (C) and (E). (**F-F"**) Anti-serotonin immunolabeling (magenta) co-labeled with 5-HT1B-MiMIC-T2A-GAL4>UAS-mCD8::GFP labeled cell bodies in the central brain. 5-HT1B-labeled kenyon cells (KC) are labeled for anatomical reference in (F"). (**G**) The approximate anatomical location for images in (F-F") are shown in the boundaries of the dashed line. Serotonin co-labeling was performed N=5 for 5-HT1B>GFP (A-A", C-C", and F-F") and N=6 brains for 5-HT1B>MCFO (E-E"). Scale bars are 20 μm.

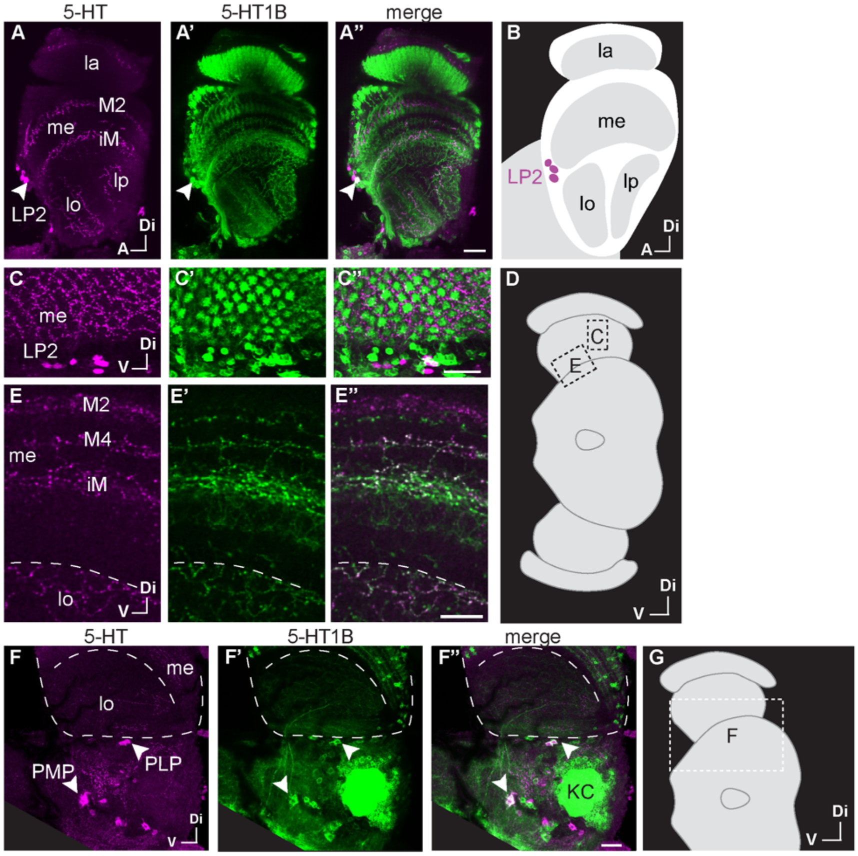

**S6 Fig. Serotonin receptor 5-HT1A co-labels with serotonin immunoreactive sites in optic lobe and cell bodies in the central brain.** (**A**) Schematic of the optic lobe neuropils—lamina (la), medulla (me), lobula (lo) and lobula plate (lp)—and serotonergic PLP cells. (**B-B"**) 5-HT1A-MiMIC-T2A-GAL4 driving UAS-mCD8::GFP (green) was co-stained with anti-serotonin immunolabeling (magenta) to map potential autoreceptors to specific cell clusters in the central brain. PLP neurons co-labeling for 5-HT1A labeling and anti-serotonin immunolabeling are indicated by the arrowhead. (**C-E**) Anterior to posterior images taken in the same brain show several serotonergic cell clusters expressing 5-HT1A (labeled at arrowheads). 5-HT1A-labeled mushroom body (MB) and kenyon cells (KC) are labeled for anatomical reference in (**C** and **E**). (**F**) A cartoon of the fly brain with dashed lines to indicate the approximate anatomical location for (C-E). Scale bars are 20 μm and N=6.

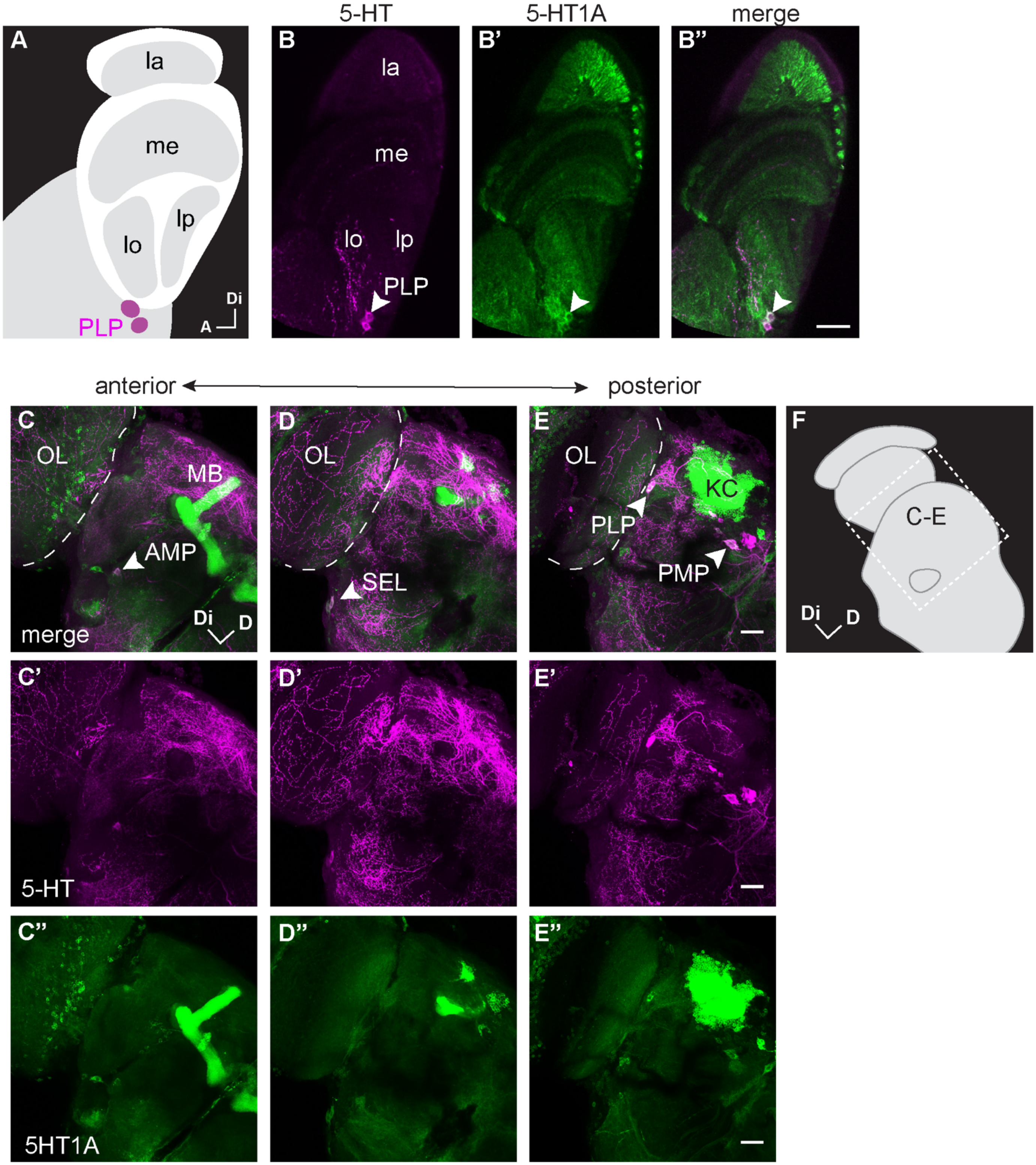
5-HT
5-HT1A
merge
A
la
me
lo
lp
PLP
Di
A
B
B'
B''
anterior
posterior
C
OL
MB
AMP
merge
Di
D
D
SEL
E
KC
PMP
F
C-E
C'
5-HT
D'
E'
C''
5HT1A
D''
E''

**S7 Fig. RT-qPCR shows L2 neurons express 5-HT2B and T1 neurons express both 5-HT1A and 5-HT1B serotonin receptors.** Enrichment (i.e., fold change) was calculated for cDNA from GFP-labeled cell isolates relative to pooled, unlabeled optic lobe cell isolates using the comparative CT method. (**A**) RT-qPCR performed on cDNA from isolated T1 neurons expressing GFP showed enrichment for serotonin receptors 5-HT1A and 5-HT1B relative to other GFP-negative cells from the optic lobe. (**B**) FACS isolates from L2 cells showed enrichment of 5-HT2B and 5-HT7 in RT-qPCR. (**C**) L1 RT-qPCR enrichment was not detectable for any serotonin receptors. RT-qPCR error bars represent mean±SEM. N=3-6 biological replicates pooled from 18-40 brains per replicate.

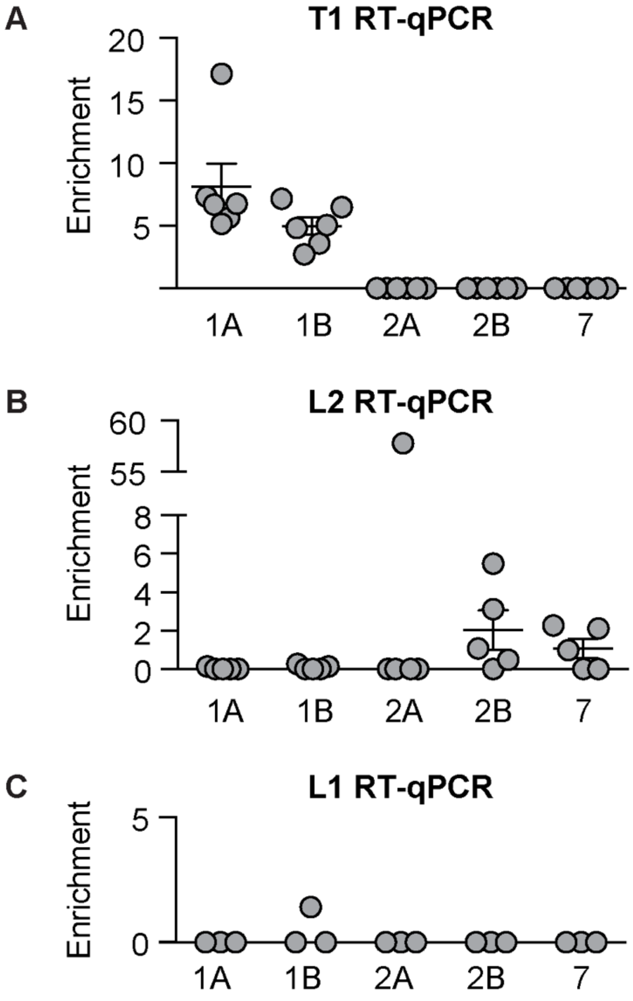

**S8 Fig**. **Serotonergic neurons do not show sybGRASP signal with postsynaptic T1, L2 or L1 neurons in the medulla.** SybGRASP was used to probe whether serotonergic neurons make synaptic contacts onto L2, T1 or L1 neurons. L2 neurons are known to synapse onto T1 projections in the medulla and we used this connection as a positive control. (**A-A'**) SybGRASP was observed with L2 split-GAL4 presynaptic to T1-LexA in M2. The dashed inset in (**A**) is shown in (**A'**). (**B-E**) A SerT-GAL4 driver was used to express the pre-synaptic portion of GFP in serotonergic neurons and LexA drivers were used to express the postsynaptic portion of GFP in L2 (**B-C**) L1 (**D**) or T1 (**E**) as indicated. No sybGRASP signal was detected in the medulla when SerT was presynaptic to L2 (**B**) however, occasional sparse GFP puncta (arrowhead) were visible in the lamina (**C**). When SerT-GAL4 neurons were presynaptic to L1 (**D**) or T1 (**E**) neurons, we did not detect a sybGRASP signal in either the lamina or medulla. All tissue was labeled with primary antibodies to both 5-HT (magenta) and GFP (green). N=7-10 brains. Scale bars are 15 µm (**A**, **B**, **D-E**); 5 µm (**A"** and **C-C"**).

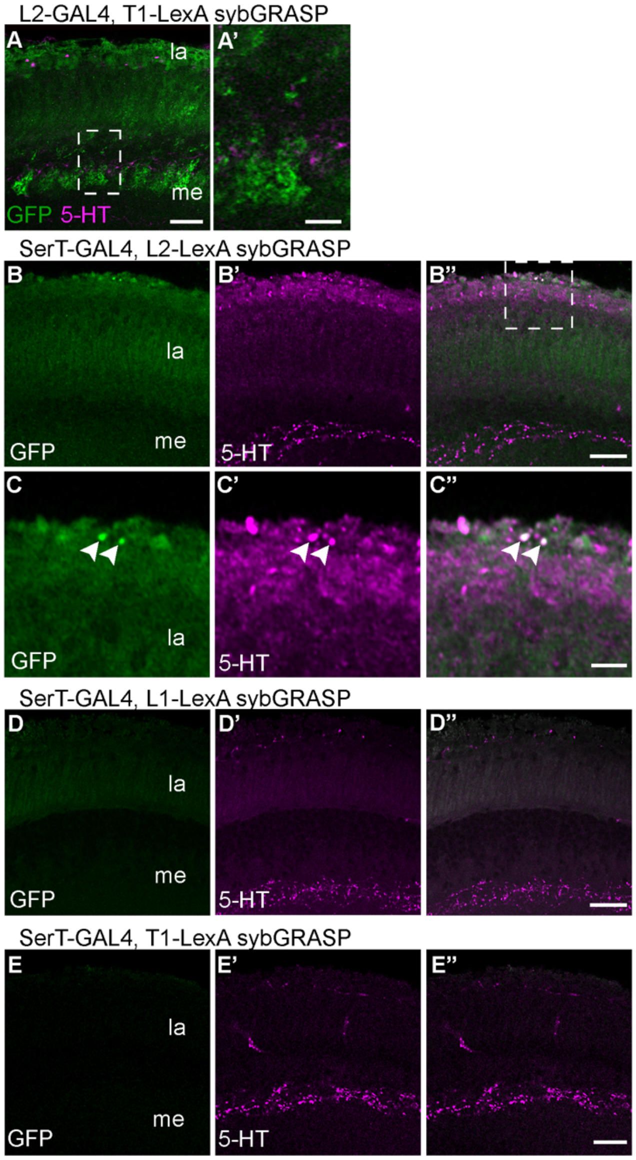

**S9 Fig. Individual traces for serotonin bath application experiments.** (**A-D**) Individual traces representing all experiments (Fig 4) for serotonin or saline controls with L2, L1 or T1 split-GAL4>GCaMP6f or T1 split-GAL4>ArcLight. For all experiments, the first 60 s of baseline is not shown; traces represent data recorded following a switch to saline with serotonin or saline alone. The length of time for the switch to complete was estimated to be 105 s, but the first 60 s of the recording are not shown so the switch occurs at 45 s on the x axis in A-D. Saline controls are gray, serotonin exposed preps are colored, and the dark line represents the mean (**A**) L2>GCaMP6f experiments, along with (**B**) L1>GCaMP6f, show an increase in calcium following serotonin application as compared to saline controls (L2, p=00095; L1, p=0.02). (**C, D**) T1 cells show no significant change with either GCaMP6f (**C**) or ArcLight (**D**) relative to saline (p>0.05) For bath application experiments (**A-D**), N=4-8 individual flies.

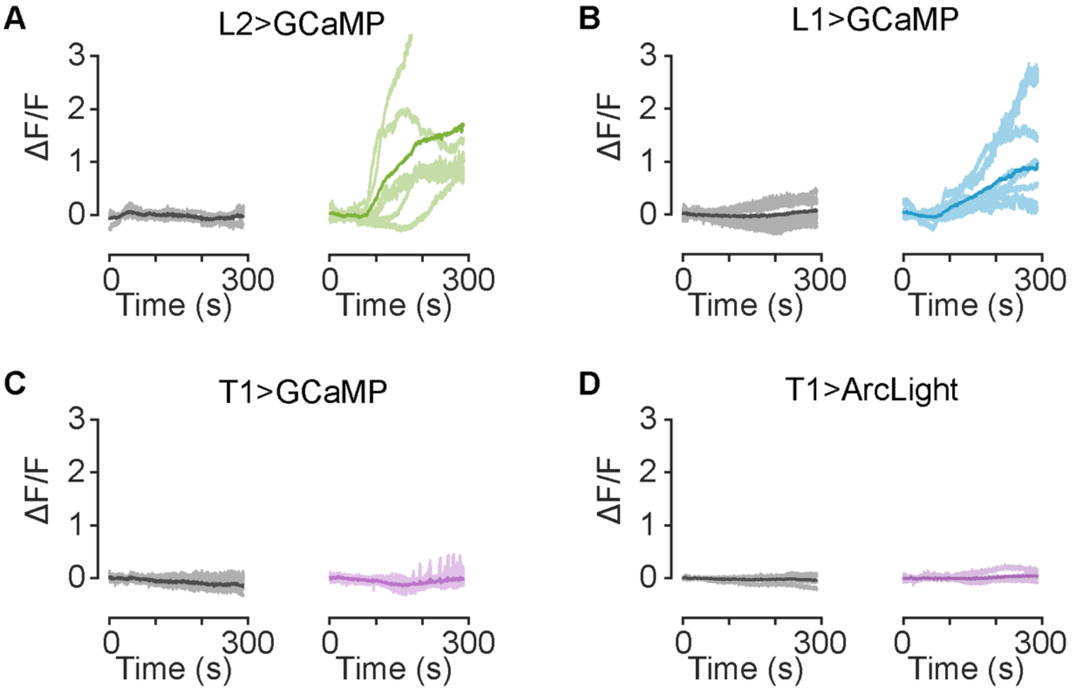

S1 Table. RNA-Seq Serotonin Receptor TPMs, averages and standard deviations.

| | 5-HT1A | 5-HT1B | 5-HT2A | 5-HT2B | 5-HT7 |
|---|---|---|---|---|---|
| T1 | 165 | 271 | 0.028 | 0.024 | 0.034 |
| T1 | 231 | 305 | 7.53 | 0.000 | 0.000 |
| T1 | 150 | 257 | 0.000 | 0.082 | 0.000 |
| Average | 182 | 278 | 2.52 | 0.035 | 0.011 |
| STDEV | 43.1 | 24.8 | 4.34 | 0.042 | 0.019 |
| | 5-HT1A | 5-HT1B | 5-HT2A | 5-HT2B | 5-HT7 |
| L2 | 0.006 | 40.9 | 13.3 | 190 | 8.96 |
| L2 | 38.3 | 22.5 | 1.13 | 45.9 | 34.2 |
| L2 | 10.4 | 28.3 | 2.99 | 158 | 3.90 |
| Average | 16.2 | 30.5 | 5.80 | 131 | 15.7 |
| STDEV | 19.8 | 9.40 | 6.55 | 75.6 | 16.2 |

S2 Table. Threshhold Cycle (CT) measurements and calculated enrichments for FACS-isolated T1, L2, and L1 samples.

| Target | T1 - Sample 1 (T1-split-GAL4) | | | T1 - Sample 2 (T1-split-GAL4) | | | T1 - Sample 3 (T1-split-GAL4) | | |
|---|---|---|---|---|---|---|---|---|---|
| | GFP+ Ave CT | GFP- Ave CT | Enrichment | GFP+ Ave CT | GFP- Ave CT | Enrichment | GFP+ Ave CT | GFP- Ave CT | Enrichment |
| RP49 | 39.7 | 32.6 | | 43.2 | 37.5 | | 38.1 | 32.4 | |
| 5HT1A | 40.3 | 36.1 | 7.29 | 43.3 | 41.7 | 17.1 | 39.4 | 36.1 | 5.64 |
| 5HT1B | 41.4 | 36.1 | 3.65 | 44.9 | 41.9 | 6.48 | 41.2 | 36.9 | 2.73 |
| 5HT2A | ND | 46.9 | | ND | ND | | ND | 47.0 | |
| 5HT2B | ND | 38.2 | | ND | 44.1 | | ND | 39.5 | |
| 5HT7 | ND | 37.1 | | | QC Failure | | ND | 37.5 | |

| Target | T1 - Sample 4 (T1-LexA) | | | T1 - Sample 5 (T1-LexA) | | | T1 - Sample 6 (T1-LexA) | | |
|---|---|---|---|---|---|---|---|---|---|
| | GFP+ Ave CT | GFP- Ave CT | Enrichment | GFP+ Ave CT | GFP- Ave CT | Enrichment | GFP+ Ave CT | GFP- Ave CT | Enrichment |
| RP49 | 37.0 | 32.5 | | 37.9 | 31.9 | | 38.6 | 33.6 | |
| 5HT1A | 39.4 | 37.3 | 5.16 | 39.9 | 36.7 | 6.70 | 39.9 | 37.7 | 6.74 |
| 5HT1B | 38.7 | 36.6 | 5.04 | 39.7 | 36.0 | 4.80 | 39.5 | 37.3 | 7.14 |
| 5HT2A | | 46.9 | | | 46.8 | | | 47.4 | |
| 5HT2B | | 39.0 | | | 38.3 | | | 39.7 | |
| 5HT7 | | 37.3 | | | 36.2 | | | 37.9 | |

| Target | L2 - Sample 1 | | | L2 - Sample 2 | | | L2 - Sample 3 | | |
|---|---|---|---|---|---|---|---|---|---|
| | GFP+ Ave CT | GFP- Ave CT | Enrichment | GFP+ Ave CT | GFP- Ave CT | Enrichment | GFP+ Ave CT | GFP- Ave CT | Enrichment |
| RP49 | 43.7 | 37.2 | | 40.9 | 35.1 | | 40.0 | 32.8 | |
| 5HT1A | ND | 42.7 | | ND | 38.2 | | 45.9 | 36.0 | 0.152 |
| 5HT1B | ND | 41.0 | | ND | 37.7 | | 45.7 | 36.6 | 0.262 |
| 5HT2A | 49.8 | 49.1 | 57.8 | ND | ND | | ND | 46.9 | |
| 5HT2B | 48.7 | 43.8 | 3.12 | 45.8 | 42.4 | 5.49 | 45.8 | 38.7 | 1.07 |
| 5HT7 | ND | 42.9 | | 44.8 | 40.0 | 2.13 | ND | 37.7 | |

| Target | L2 - Sample 4 | | | L2 - Sample 5 | | |
|---|---|---|---|---|---|---|
| | GFP+ Ave CT | GFP- Ave CT | Enrichment | GFP+ Ave CT | GFP- Ave CT | Enrichment |
| RP492 | 40.1 | 34.1 | | 38.4 | 34.3 | |
| 5HT1A | ND | 35.9 | | ND | 40.2 | |
| 5HT1B | 45.2 | 36.4 | 0.151 | ND | 40.1 | |
| 5HT2A | ND | ND | | ND | ND | |
| 5HT2B | 44.8 | 37.7 | 0.495 | ND | 43.3 | |
| 5HT7 | 41.6 | 36.7 | 2.266 | 45.0 | 40.8 | 0.99 |

| Target | L1 - Sample 1 | | | L1 - Sample 2 | | | L1 - Sample 3 | | |
|---|---|---|---|---|---|---|---|---|---|
| | GFP+ Ave CT | GFP- Ave CT | Enrichment | GFP+ Ave CT | GFP- Ave CT | Enrichment | GFP+ Ave CT | GFP- Ave CT | Enrichment |
| RP49 | 45.3 | 36.7 | | 42.4 | 38.0 | | 41.9 | 32.8 | |
| 5HT1A | ND | 36.4 | | ND | 41.5 | | ND | 37.5 | |
| 5HT1B | ND | 36.8 | | ND | 41.9 | | 45.7 | 37.1 | 1.416 |
| 5HT2A | ND | 41.2 | | ND | ND | | ND | ND | |
| 5HT2B | ND | 40.4 | | ND | 43.6 | | ND | 40.6 | |
| 5HT7 | ND | 37.5 | | ND | 45.1 | | ND | 39.1 | |

S3 Table. Fly Strains used in this study.

| Drivers | Source | Figure |
|---|---|---|
| 5-HT1A-T2A-GAL4 MI01468 | H. Dierick (Baylor) | S1B, S3E, S3D, S6B-E |
| 5-HT1A-T2A-GAL4 MI01140 | H. Dierick (Baylor) | 1A |
| 5-HT1A-T2A-GAL4 MI04464 | H. Dierick (Baylor) | S3G |
| 5-HT1B-T2A-GAL4 MI05213 | H. Dierick (Baylor) | 1B, S1C, S5A-F |
| 5-HT2A-T2A-GAL4 MI0459 | H. Dierick (Baylor) | S1C, S4B |
| 5-HT2A-GAL4 MI03299 | H. Dierick (Baylor) | S4A |
| 5-HT2B-T2A-GAL4 MI5208 | H. Dierick (Baylor) | 1D, S1E, S3A |
| 5-HT7-GAL4 MI00215 | H. Dierick (Baylor) | 1E, S1F, S3B-C, S3F, S3H |
| T1-spGAL4 | A. Nern (Janelia) | 4D, S7A, S9C-D |
| L2-spGAL4 | L. Zipursky (UCLA) | 2B, 3C-D, 4B, 5A-E, 6A, S7B, S8A, S9A |
| L1-spGAL4 | A. Nern (Janelia) | 4C, S7C, S9B |
| T1-LexA | A. Nern (Janelia) | 2A, S7A, S8A, S8E |
| L2-LexA | RRID:BDSC_52510 | S8B-C |
| L1-LexA | A. Nern (Janelia) | S8D |
| SerT-Gal4 (P{GMR50H05-GAL4}attP2) | RRID:BDSC_38764 | S8B-E |
| Chat-LexA MI | RRID:BDSC_60319 | S3A |
| GAD1-LexA | RRID:BDSC_60324 | S3E |

| Reporters and Mutants | Source | Figure |
|---|---|---|
| UAS-mCD8::GFP | RRID:BDSC_5137 | 2A-B, 3D, S4B, S5A, S5C-F, S6B-E, S7A-C |
| UAS-mCD8::RFP, LexAop-mCD8::GFP (x) | RRID:BDSC_32229 | S1B-F, S3A, S3E |
| UAS-MCFO-1 | RRID:BDSC_64085 | 1A-E, S3B-D, S3F-H, S4A, S5C |
| UAS-GCaMP6f | RRID:BDSC_42747 | 4B-D, 5A-E, 6A-F, S9A-C |
| UAS-ArcLight | RRID:BDSC_51056 | S9D |
| (sybGRASP) UAS-nSyb::GFP1-10, LexAop-CD4:GFP11 | RRID:BDSC_64314 | S8A-E |
| 5-HT1A::GFP | Y. Rao (Peking U.) | 3A-B |
| UAS-5-HT2B::GFP | Y. Rao (Peking U.) | 3C |
| 5-HT2B-GKO-GAL4 | Y. Rao (Peking U.) | 6B-F |

S4 Table. Primer sequences and mRNA target information.

| Primer ID | Gene Symbol | Exon Junction | Product Length (bp) | Primer Sequence (5' to 3') | Primer Length | One Primer Spans | Transcript Specificit | Design |
|---|---|---|---|---|---|---|---|---|
| 5HT1A Forward | Dmel\5-HT1A | 2027/2028 | 159 | TTCGTGGCCTGCCTAGTAAT | 20 | Yes | All | BLAST |
| 5HT1A Reverse | Dmel\5-HT1A | 2027/2028 | 159 | CCAGTAACGATCGACGGCAA | 20 | Yes | All | BLAST |
| 5HT2B Forward | Dmel\5-HT2B | 302/303 | 135 | AAAGCCGATTGCTTCTCCAAC | 21 | Yes | All | BLAST |
| 5HT2B Reverse | Dmel\5-HT2B | 302/303 | 135 | GATTCAGGACTCGCGAAAGG | 20 | Yes | All | BLAST |
| 5HT1B Forward | Dmel\5-HT1B | 2148/2149 | 129 | ATTTCGCCAGTTTGGCCATT | 20 | Yes | All | BLAST |
| 5HT1B Reverse | Dmel\5-HT1B | 2148/2149 | 129 | CGTTGCTGGTGCGATAATCA | 20 | Yes | All | BLAST |
| 5HT7 Forward | Dmel\5-HT7 | 700/701 | 117 | TCGACGACTTTTGGAAGCAC | 20 | Yes | All | BLAST |
| 5HT7 Reverse | Dmel\5-HT7 | 700/701 | 117 | ATTGTCGTCGGGAACTGGG | 19 | Yes | All | BLAST |
| 5HT2A Forward | Dmel\5-HT2A | 480/481 | 186 | CGACTGCAAAATGTGGGTCT | 20 | No | All except F | BLAST |
| 5HT2A Reverse | Dmel\5-HT2A | 480/481 | 186 | CCTCGCAGATGTGTCTGTTG | 20 | No | All except F | BLAST |
| RP49 Forward | Dmel\RpL32 | 28/29 | 147 | GGTTTCCGGCAAGCTTCAA | 19 | Yes | B Only | BLAST |
| RP49 Reverse | Dmel\RpL32 | 28/29 | 147 | TGTTGTCGATACCCTTGGGC | 20 | Yes | B Ony | BLAST |
